# Differential spreading behaviors of Nodal signalling molecules in the extracellular space cooperatively shape left-right asymmetry

**DOI:** 10.1101/2025.08.13.670121

**Authors:** Takafumi Ikeda, Toru Kawanishi, Yusuke Mii, Taijiro Yabe, Jose Pelliccia, Rebecca D Burdine, Hiroyuki Takeda

## Abstract

Left-right (LR) patterning of vertebrate embryos is mediated by diffusive proteins secreted mainly from the left-right organizer (LRO), such as TGF-β superfamily ligands Nodal and Gdf1, as well as their inhibitors Dand5 and Lefty. However, precise modes of action for these proteins remain unclear due to difficulties in visualizing and manipulating their extracellular behaviors. Here we visualize and manipulate the extracellular dynamics of these proteins employing novel transgenic zebrafish lines and extracellular trapping molecules including the morphotrap. We demonstrate that the heterodimer of Nodal and Gdf1, which is the major signal transducer in vertebrate Nodal signalling, spreads from the LRO through the extracellular space in the paraxial mesoderm (PM) to its target tissue, the lateral plate mesoderm (LPM), in a free-diffusion-like manner. By contrast, Dand5 is highly enriched in the right PM, where it is immobilized through interaction with heparan sulfate proteoglycans and asymmetrically suppresses Nodal signalling. Our study uncovers diverse modes of action for LR-related secreted proteins in the extracellular space, providing a new framework for understanding how spatially organized Nodal signalling molecules shape embryonic LR asymmetry.

## Introduction

Embryonic pattern formation is regulated by various signalling ligands and inhibitor proteins secreted from source cells. Signalling ligands spread through the extracellular space in the embryo and induce specific gene expression in target tissues, typically in a concentration-dependent manner^1–3^. Secreted inhibitor proteins also spread within the extracellular space and modulate the signalling activity and range of their target ligands^4^. While genetic analyses have revealed their broad functions in embryo patterning, protein-level analyses are essential for understanding their mechanisms of action, including ligand distribution range, secretion and internalization rate, and affinity for the extracellular matrix (ECM). Recent advances in fluorescent imaging have enabled us to visualize the extracellular dynamics of signalling ligands in certain model systems, such as the wing imaginal disc of *Drosophila* and the zebrafish blastula^5–11^. However, it is still challenging to know how the signalling ligands and inhibitors behave in the extracellular space of vertebrate embryos, especially at later stages, mainly due to their low extracellular concentrations and the increased structural complexity of the embryo. Left-right (LR) asymmetry formation in vertebrate embryos provides a fascinating yet complicated model for studying the behavior of signalling ligands and their inhibitors. LR asymmetry is formed through the following steps: emergence and sensing of the leftward fluid flow in the left-right organizer (LRO), signal transmission from the LRO to the lateral plate mesoderm (LPM), and asymmetric visceral organ formation driven by the LPM^12,13^. The initial symmetry-breaking event occurs in the LRO (i.e., the mouse node and teleost Kupffer’s vesicle (KV)), where unidirectional rotation of motile cilia produces the leftward fluid flow and evokes asymmetric Ca^2+^ signalling in the surrounding epithelium^12,13^. LRO cells secrete a TGF-β superfamily ligand Nodal and a DAN-family inhibitor Dand5, both of which are required for proper LR patterning^9,14,15^. Secreted Nodal is thought to transmit its signal to the left LPM, and induce its own expression^16^. Nodal is likely to form a heterodimer in the LRO with another TGF-β superfamily ligand, Gdf1 (also known as Gdf1/3 in bony fish and Vg1 in anurans^17^), which is also required for the left-sided *nodal* expression in the LPM^9,18,19^. While *nodal* and *gdf1* are bilaterally expressed, *dand5* is expressed preferentially on the right side of the LRO, and suppresses Nodal signalling on the right^14,20^. Furthermore, another Nodal inhibitor, Lefty, is expressed in the midline tissues (and in the left LPM in mouse embryos) and restricts Nodal activity in the left LPM^21,22^. Together, LR asymmetry formation in the LPM is regulated by the coordinated actions of the Nodal-Gdf1 ligand and its inhibitors, Dand5 and Lefty (**Fig. 1a**).

**Figure 1.**
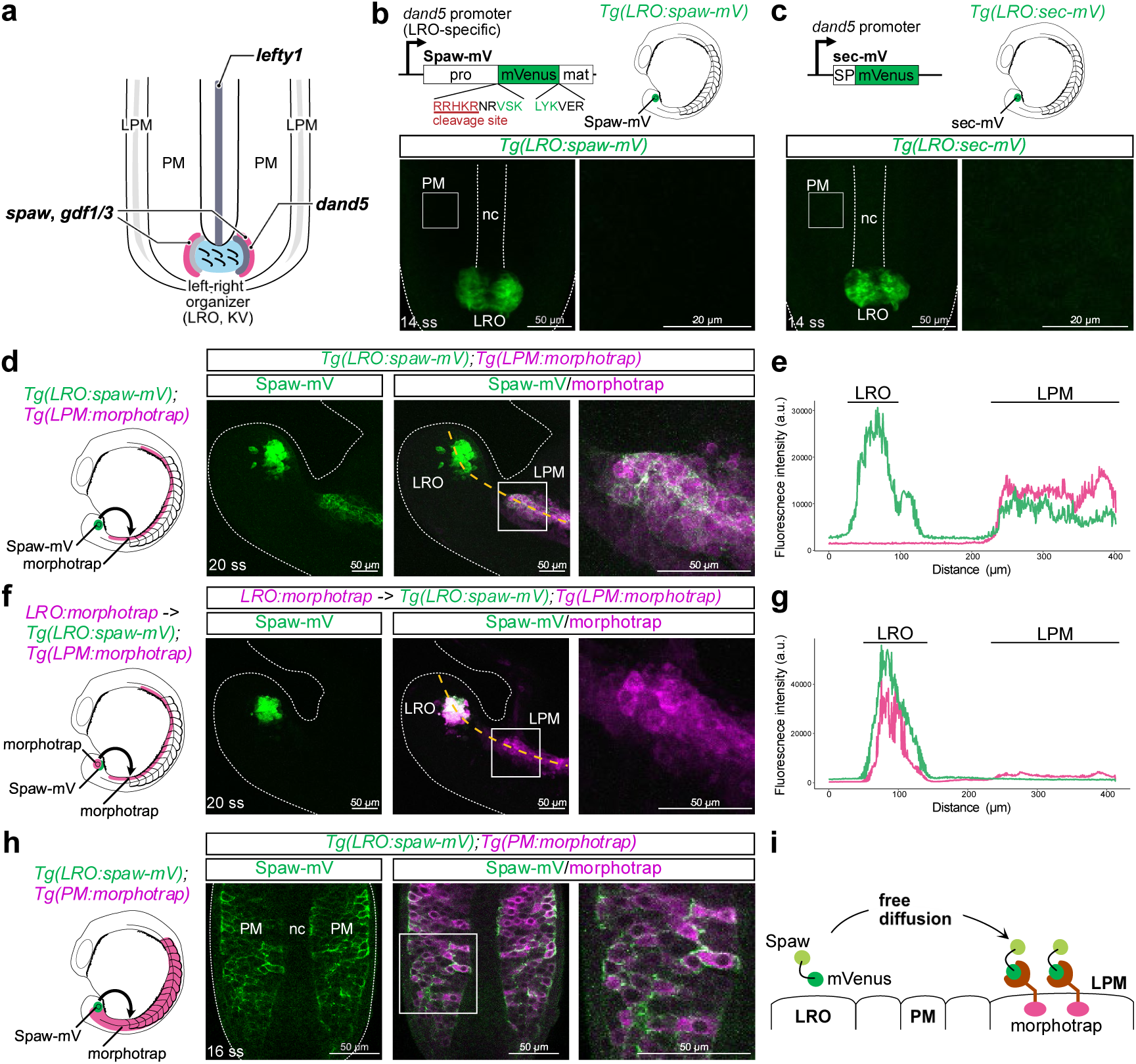
Extracellular distribution of Spaw. **a,** Expression of *spaw*, *gdf1/3*, *dand*5 and *lefty1* in the zebrafish embryo at early segmentation stage (i.e., LR asymmetry formation period). **b, c,** Extracellular distribution of Spaw-mV (b) and sec-mV (c) in 14-ss *Tg(LRO:spaw-mV)* (b) and *Tg(LRO:sec-mV)* (c) embryos. Dorsal view of the tailbud. mVenus CDS is inserted two amino acids downstream of the prodomain cleavage site of *spaw*. nc, notochord. pro, prodomain. mat, mature domain. SP, signal peptide. **d,** Double transgenic line, *Tg(LRO:spaw-mV);Tg(LPM:morphotrap)*. Lateral view of the tailbud. **e,** Fluorescence intensity of Spaw-mV and morphotrap along the anterior-posterior axis from the LRO to the LPM (yellow line in d). Green, Spaw-mV. Magenta, morphotrap. **f,** Suppressed diffusion of Spaw-mV by the morphotrap expression. *LRO:morphotrap* plasmid was injected into *Tg(LRO:spaw-mV);Tg(LPM:morphotrap)* embryos at 1-cell stage. **g,** Fluorescence intensity of Spaw-mV and morphotrap along the anterior-posterior axis from the LRO to the LPM (yellow line in f). Green, Spaw-mV. Magenta, morphotrap. **h,** Double transgenic line, *Tg(LRO:spaw-mV);Tg(PM:morphotrap)*. White dotted lines, contour of embryos. White boxes, magnified areas. **i,** Schematic of Spaw diffusion from the LRO to the LPM through the extracellular space of the PM. Maximum intensity projection is shown for (b), (c), (d) and (f). *n =* 13 (b), 12 (c), 12 (d), 8 (f), 4 (h) embryos.

Despite their importance, as supported by extensive genetic analyses, few studies have investigated the behavior of Nodal-Gdf1, Dand5 and Lefty proteins in the extracellular space during LR asymmetry formation. These proteins are also expressed prior to LR asymmetry formation and involved in germ layer patterning^22,23^. Quantitative measurements of labelled Nodal and Lefty proteins, overexpressed in the zebrafish blastula by mRNA injection, demonstrated that they form a reaction-diffusion system^6,7,24^. However, mRNA injection is not suitable for investigating the behavior of Nodal-Gdf1, Dand5, and Lefty during LR asymmetry formation, as their expression at this stage is spatially restricted and overexpression of these proteins can disrupt earlier developmental events, including germ layer formation. Consequently, how Nodal-Gdf1 heterodimers secreted from the LRO behave in the extracellular space to transmit signals to the distant LPM remains elusive. Similarly, it is unclear how the inhibitor proteins Dand5 and Lefty exhibit distinct extracellular behavior to spatially restrict Nodal signalling. Addressing these questions requires direct visualization and manipulation of these proteins under physiologically relevant conditions, which has not been achieved to date.

Utilizing transgenic technology in zebrafish, we developed an *in vivo* reconstitutive system for imaging fluorescently labelled LR-related ligands and inhibitors. This system is designed to mimic their endogenous expression patterns while enhancing expression levels to facilitate confocal imaging. By combining this system with ligand-trapping technology, we visualized their extracellular behaviors of these molecules and manipulated their activity during LR patterning. These analyses revealed the differential modes of action for Nodal-Gdf1 and its inhibitors: Nodal-Gdf1 and Lefty exhibit free-diffusion-like behavior in the paraxial mesoderm (PM) located between the LRO and LPM, whereas Dand5 is strongly associated with heparan sulfate proteoglycans (HSPGs) in the right PM, where it locally inhibits Nodal signal transmission to the LPM. Our findings demonstrate the asymmetric and cooperative interplay between signalling ligands and inhibitors in the extracellular space, which is essential for proper LR asymmetry formation in the LPM.

## Results

### Extracellular distribution of Spaw

To visualize the distribution of Nodal proteins secreted from the LRO, we expressed mVenus (mV)-tagged Spaw, a zebrafish Nodal homolog involved in LR patterning^15^, in the LRO. Unlike in mouse embryos, teleost *spaw* is expressed beside KV (the LRO in teleosts) but not in KV itself^15,25^. However, since the regulatory sequence that reproduces endogenous *spaw* expression has not been identified yet, we instead used the KV-specific *dand5* promoter^26^ to express Spaw-mV in KV-epithelial cells (*Tg(LRO:spaw-mV)*; **Fig. 1b**). For comparison, we also made *Tg(LRO:sec-mV)*, which expresses a secreted form of mVenus in the LRO (**Fig. 1c**). Confocal microscopy of the *Tg(LRO:spaw-mV)* and *Tg(LRO:sec-mV)* embryos immunostained with an anti-GFP antibody (which recognizes mV) revealed that the signals of Spaw-mV and sec-mV were detected only in the LRO, but not in adjacent tissues (**Fig. 1b, c**). This suggests that the extracellular Spaw-mV proteins were below the detection threshold for confocal microscopy. To circumvent this limitation and enhance detection sensitivity, we introduced the morphotrap, a membrane-tethered GFP nanobody tagged with an intracellular mCherry domain, which improves the detection sensitivity of diffusing GFP (or its derivatives)-tagged proteins by accumulating them on the cell surface^5,27–29^ (**Fig. 1i**). We established *Tg(LPM:morphotrap)*, a transgenic line expressing the morphotrap specifically in the LPM under the control of the LPM-specific *drl* promoter^30^ (**Fig. 1d**). Remarkably, in the double transgenic *Tg(LRO:spaw-mV);Tg(LPM:morphotrap)* embryos, the signal of Spaw-mV was detected on the cell surface of the LPM (**Fig. 1d, e; Supplementary Fig. 1**), suggesting that the morphotrap expressed in the LPM cells traps extracellular Spaw-mV. To corroborate that Spaw-mV trapped in the LPM is directly derived from the LRO, we expressed the morphotrap in the LRO (the source of Spaw-mV) of *Tg(LRO:spaw-mV);Tg(LPM:morphotrap)* embryos by injecting *LRO:morphotrap* plasmid. The signal of Spaw-mV in the LPM was greatly diminished in injected embryos that uniformly expressed the morphotrap in the LRO (**Fig. 1f, g**), confirming that Spaw-mV spreads from the LRO to the LPM. Spaw-mV signal in *Tg(LRO:spaw-mV);Tg(LPM:morphotrap)* embryos was detected bilaterally at the posterior part of the LPM from 16-somite stage (ss) and gradually extended anteriorly (**Supplementary Fig. 1**). Furthermore, in *Tg(LRO:HA-spaw-mV);Tg(LPM:morphotrap)* embryos, in which 4×HA tags were inserted into the N-terminus of the prodomain of Spaw-mV, the mature domain of HA-Spaw-mV was detected only in the LPM, while the HA-tagged prodomain was confined to the LRO (**Supplementary Fig. 2**). This suggests that Spaw proteins reaching the LPM lack their prodomains, possibly due to intracellular cleavage, as observed in other TGF-β superfamily ligands^31^.

We then crossed *Tg(LRO:spaw-mV)* with *Tg(PM:morphotrap)*, in which the morphotrap is specifically expressed in the PM, the intermediate tissue located between the LRO and LPM, under the control of the PM-specific *msgn1* promoter^32^. We found that Spaw-mV was accumulated on the morphotrap-expressing PM cells (**Fig. 1h; Supplementary Fig. 3**), supporting the idea that Spaw-mV secreted from the LRO spreads through the extracellular space within the PM to the LPM (**Fig. 1i**). By contrast, when the morphotrap was expressed specifically in the hatching gland (HG), which is located far from the LRO, using of the HG-specific *he1.1* promoter^33^, Spaw-mV did not accumulate on the morphotrap-expressing HG cells (**Supplementary Fig. 4**). These results indicate that the morphotrap can detect Spaw-mV only when it is present at sufficiently high levels in the embryo. They also suggest that Spaw-mV is specifically distributed in the posterior region near the LRO, rather than spreading throughout the entire embryo.

### Extracellular distribution of Gdf1/3 and Spaw-Gdf1/3 heterodimer

We next analyzed the distribution of Gdf1/3 (a zebrafish ortholog of Gdf1^17^), which acts as a co-ligand of Nodal, by establishing *Tg(LRO:gdf1/3-mV)* (**Fig. 2a**). Contrary to the *Tg(LRO:spaw-mV);Tg(LPM:morphotrap)* embryos, the *Tg(LRO:gdf1/3-mV);Tg(LPM:morphotrap)* embryos showed no accumulation of Gdf1/3-mV in the LPM, suggesting that Gdf1/3-mV hardly diffuses to the LPM (**Fig. 2b, c**), consistent with the observation in the zebrafish blastula that Gdf1/3 tends to remain in source cells in the absence of Nodal^9^.

**Figure 2.**
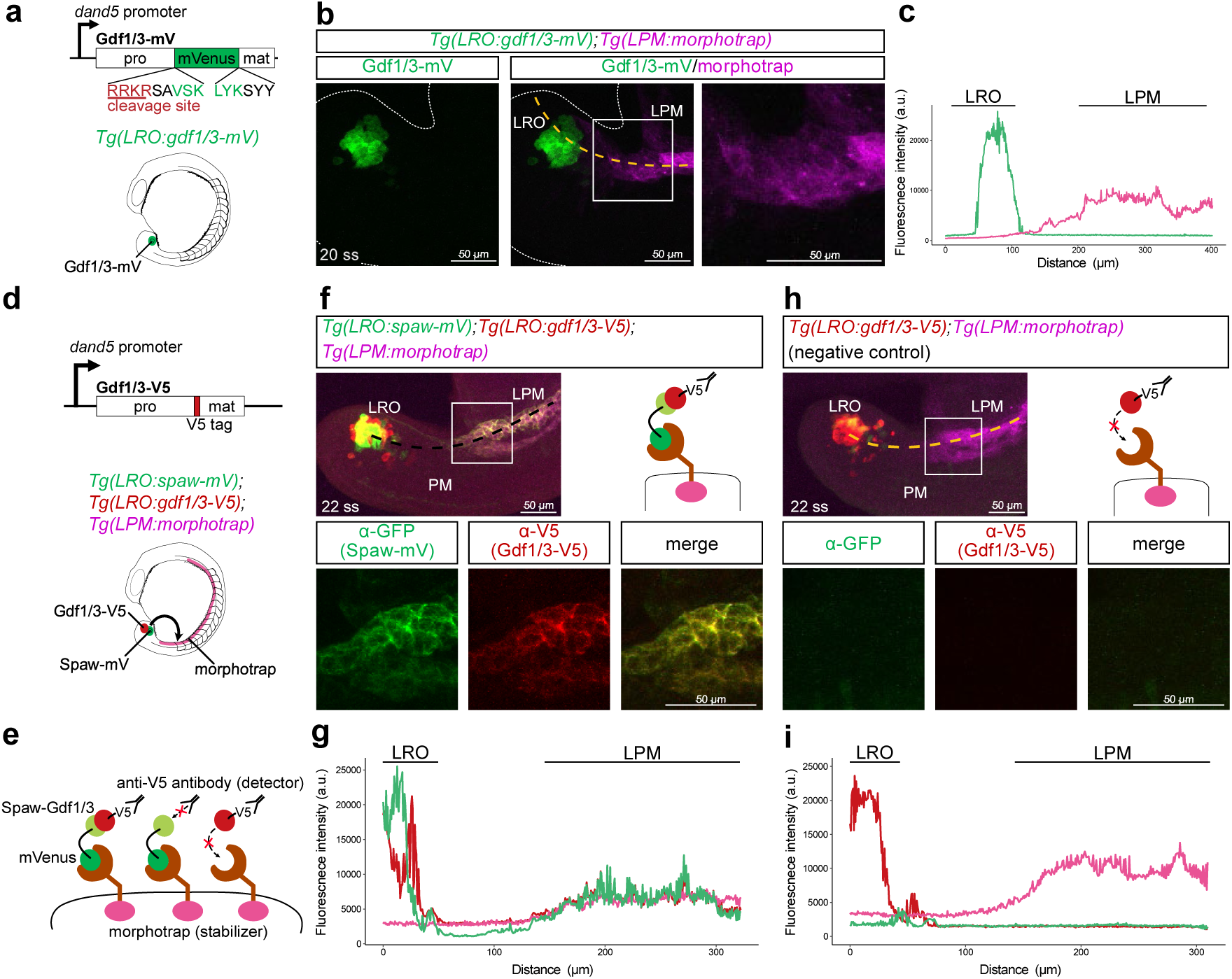
Detection of Spaw-Gdf1/3 heterodimer by *in situ* sandwich assay. **a,** *Tg(LRO:gdf1/3-mV)*. mVenus CDS is inserted two amino acids downstream of the prodomain cleavage site. pro, prodomain. mat, mature domain. **b,** Extracellular distribution of Gdf1/3-mV in *Tg(LRO:gdf1/3-mV);Tg(LPM:morphotrap)*. White dotted lines, contour of embryos. White boxes, magnified areas. **c,** Fluorescence intensity of Gdf1/3-mV and morphotrap along the anterior-posterior axis from the LRO to the LPM (yellow line in b). Green, Gdf1/3-mV. Magenta, morphotrap. **d,** Triple transgenic line, *Tg(LRO:spaw-mV);Tg(LRO:gdf1/3-V5);Tg(LPM:morphotrap)*. Gdf1/3-V5 has a V5 tag in the N-terminal of Gdf1/3 mature domain. **e,** Schematic of *in situ* sandwich assay. The morphotrap on the LPM cells stabilizes both Spaw-mV alone and Spaw-mV-Gdf1/3-V5, but not Gdf1/3-V5 alone. By immunostaining with anti-V5 antibody, Spaw-mV-Gdf1/3-V5 but not Spaw-mV alone is detected. **f-i,** Detection of Spaw-mV-Gdf1/3-V5 heterodimer in the LPM. 20-ss embryos of *LRO:spaw-mV;LRO:gdf1/3-V5;LPM:morphotrap* (f) and *LRO:gdf1/3-V5;LPM:morphotrap* (h) were immunostained with anti-GFP and anti-V5 antibodies. Lateral view of the tailbud. White boxes, magnified areas. Fluorescence intensity of Spaw-mV, Gdf1/3-V5 and morphotrap along the black (f) and yellow (h) lines was plotted in (g) and (i), respectively. Green, Spaw-mV. Red, Gdf1/3-V5. Magenta, morphotrap. Maximum intensity projections of each image are shown. *n =* 6 (b), 7 (f), 3 (h) embryos.

Previous studies have suggested that the Nodal-Gdf1 heterodimer, which exerts a longer-range effect than Nodal alone, is a major ligand in vertebrate Nodal signalling^8,18,34,35^. In *Tg(LRO:spaw-mV);Tg(LPM:morphotrap)* used above, it was unclear whether Spaw-mV that was trapped by the morphotrap in the LPM was a homodimer (or a monomer) or a heterodimer with endogenous Gdf1/3, although the endogenous expression level of Gdf1/3 was expected to be much lower than that of transgenic Spaw-mV. To specifically visualize the diffusion of Spaw-Gdf1/3 heterodimers in the embryo, we introduced Gdf1/3 tagged with the V5 epitope (Gdf1/3-V5) into our reconstitutive system by generating a triple transgenic line, *Tg(LRO:spaw-mV); Tg(LRO:gdf1/3-V5); Tg(LPM:morphotrap)* (**Fig. 2d**). In this line, Spaw-mV and Gdf1/3-V5 were co-expressed in the LRO, while the morphotrap was expressed in the LPM. We then performed immunostaining to detect both Spaw-mV and Gdf1/3-V5. The *Tg(LRO:gdf1/3-V5);Tg(LPM:morphotrap)* embryos were similarly processed as a negative control. This “*in situ* sandwich” assay would allow specific detection of epitope-tagged Spaw-Gdf1/3 heterodimers in embryos^10^ (**Fig. 2e**). In *Tg(LRO:spaw-mV);Tg(LRO:gdf1/3-V5);Tg(LPM:morphotrap)* embryos, Spaw-mV and Gdf1/3-V5 were specifically colocalized around the LRO and on LPM cells, but not on PM cells located between them (**Fig. 2f, g**). Their signals were undetectable in the LPM of *Tg(LRO:gdf1/3-V5);Tg(LPM:morphotrap)* embryos (**Fig. 2h, i**). This result indicates that Spaw and Gdf1/3 diffuse to the LPM as a heterodimer. By contrast, no anti-Spaw-V5 signal was detected in the LPM of the *Tg(LRO:spaw-mV);Tg(LRO:spaw-V5);Tg(LPM:morphotrap)* embryos (**Supplementary Fig. 5**), suggesting that Spaw alone hardly forms a homodimer that effectively diffuses to the LPM. Altogether, these findings demonstrate that the Nodal-Gdf1 heterodimer, but not Nodal homodimer, spreads from the LRO to the LPM during LR asymmetry formation.

### Extracellular distribution of Dand5 and Lefty1

We next visualized the distribution of Dand5, a right-side Nodal inhibitor secreted from the LRO, by establishing *Tg(LRO:dand5-mV)* (**Fig. 3a**). In sharp contrast to Spaw-mV, Gdf1/3-mV and sec-mV, Dand5-mV was detected outside the LRO in *Tg(LRO:dand5-mV)* embryos by immunostaining without morphotrap expression. It accumulated in the PM in a punctate manner at 14 ss and weakly detected around the notochord at 20 ss, when *spaw* expression in the left LPM is established (**Fig. 3b; Supplementary Fig. 6**). Those puncta were mainly distributed in the extracellular space, but were also observed intracellularly, suggesting that they were internalized into PM cells (**Fig. 3b**). Similar distribution was observed in *Tg(LRO:dand5-OLLAS)* embryos, in which Dand5 tagged with the OLLAS epitope is expressed in the LRO (**Supplementary Fig. 7a**). Furthermore, in the *Tg(LRO:dand5-mEosEM)*, which expresses Dand5 tagged with mEosEM, a bright photoconvertible protein^36^, Dand5-mEosEM similarly showed punctate localization in the PM of live embryos, confirming that the observed puncta are not fixation artifacts (**Supplementary Fig. 7b**).

**Figure 3.**
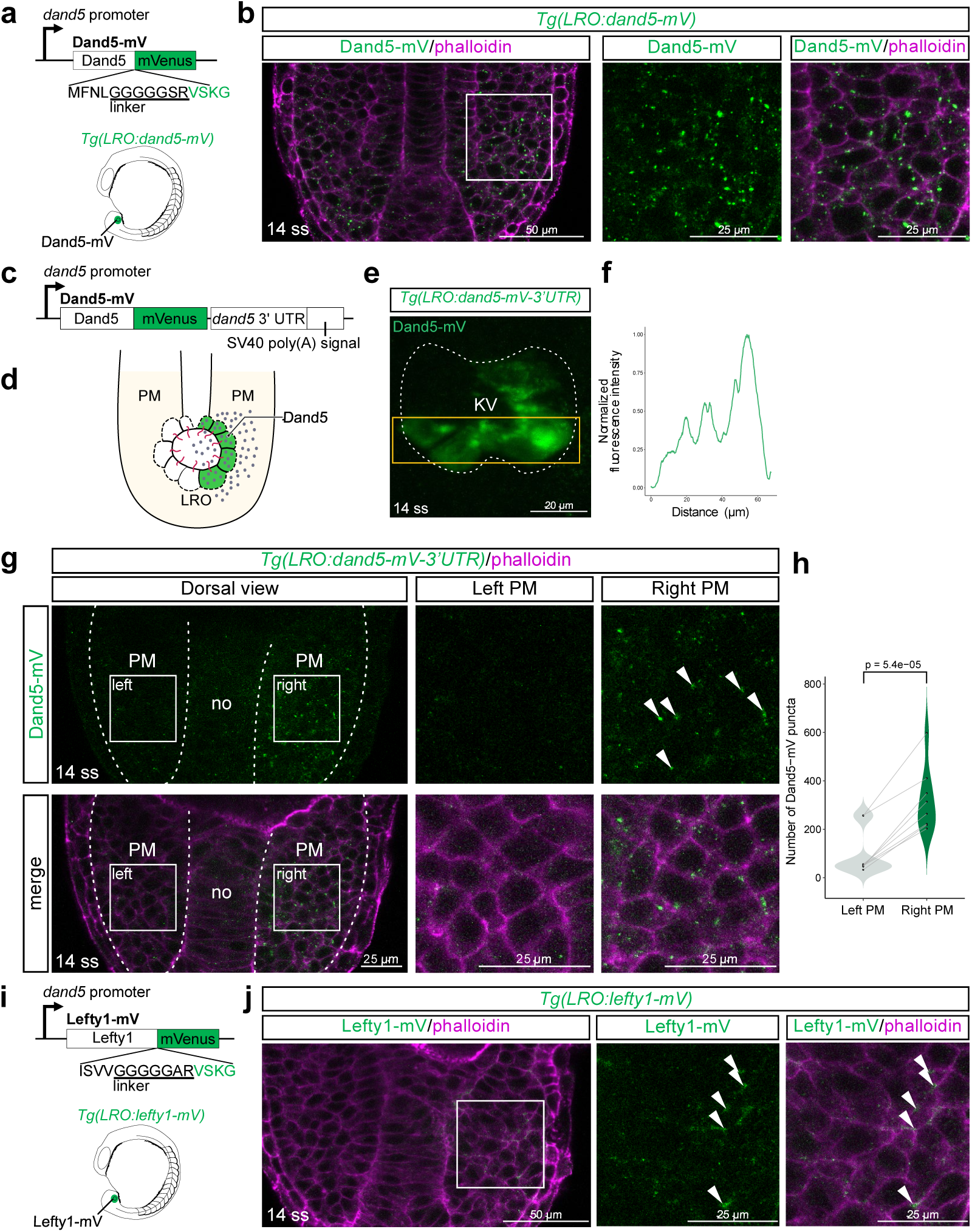
Extracellular distribution of Dand5 and Lefty. **a,** *Tg(LRO:dand5-mV)*. mVenus is fused to the C-terminus of Dand5 or Lefty1 with 5× glycine linker. **b,** Extracellular distribution of Dand5-mV in *Tg(LRO:dand5-mV)*. Embryos were counterstained with phalloidin to highlight cell membranes. Dorsal view of the tailbud. **c,** *Tg(LRO:dand5-mV-3’UTR)*. 3’ UTR sequence of *dand5* is inserted downstream of CDS for Dand5-mV, which is followed by SV40 poly(A) signal to enhance expression level. **d,** Schematic of the distribution pattern of Dand5 secreted from the LRO. **e,** Distribution of Dand5-mV in the LRO of a *Tg(LRO:dand5-mV-3’UTR)* embryo. Dorsal view of the tailbud. Maximum intensity projection is shown. **f,** Normalized fluorescence intensity plot for (e). Fluorescence intensity in the yellow box in (e) was measured along the left-right axis. **g, h,** Asymmetric distribution of Dand5-mV in the PM of *Tg(LRO:dand5-mV-3’UTR)* embryo. Embryos were counterstained with phalloidin to highlight cell membranes. Dorsal view of the tailbud (g). White arrowheads, puncta of Dand5-mV. White boxes, magnified areas. The number of Dand5-mV puncta in the left and right PM was quantified in (h) (see M&M for detail). *p*-value was calculated using a paired *t*-test. **i,** *Tg(LRO:lefty1-mV)*. mVenus is fused to the C-terminus of Lefty1 with 5× glycine linker. **j,** Extracellular distribution of Lefty1-mV in *Tg(LRO:lefty1-mV)*. Embryos were counterstained with phalloidin to highlight cell membranes. Dorsal view of the tailbud. *n* = 10 (b), 5 (e), 8 (g, h), 9 (j) embryos.

In the *Tg(LRO:dand5-mV)* embryos, Dand5-mV was symmetrically expressed in the LRO and distributed in the PM, which did not recapitulate the right-sided endogenous expression of *dand5*. Previous studies reported that the 3’ untranslated region (UTR) of *dand5* mRNA is responsible for left-sided degradation of *dand5* mRNA in mouse and *Xenopus* embryos^37,38^. We thus cloned the 3’ UTR sequence from zebrafish *dand5* mRNA and inserted it into the transgene cassette to establish *Tg(LRO:dand5-mV-3’UTR)* (**Fig. 3c**). In *Tg(LRO:dand5-mV-3’UTR)* embryos, Dand5-mV was accumulated preferentially on the right side of the LRO (**Fig. 3d-f**), indicating the essential role of the 3’UTR sequence of zebrafish *dand5* mRNA for its right-sided expression. Importantly, Dand5-mV puncta were maintained on the right side of the PM, but were almost absent on the left side in *LRO:dand5-mV-3’UTR* embryos (**Fig. 3d, g, h**). Together, we concluded that Dand5 secreted from the right side of the LRO is asymmetrically distributed in the extracellular space of the right PM in a punctate manner.

We next compared the distribution pattern of Dand5 with that of the other LR-related Nodal inhibitor Lefty (Lefty1), which is expressed in the midline tissue of zebrafish embryos^22^. For direct comparison, we expressed Lefty1-mV in the LRO cells using the *dand5* promoter (**Fig. 3i**). Unlike Dand5-mV in *Tg(LRO:dand5-mV)* embryos, Lefty1-mV hardly showed punctate localization in the PM of *Tg(LRO:lefty1-mV)* embryos, but instead weakly accumulated in the tricellular space within the PM and the space between the PM and the ectoderm (**Fig. 3j**). These results demonstrate that the two Nodal inhibitors exhibit distinct extracellular behaviors in embryos.

### Heparan sulfate proteoglycans are involved in Dand5 punctate distribution

The punctate distribution of Dand5-mV in the PM suggests that Dand5 is associated with interacting molecules in the extracellular space, most likely with extracellular matrix (ECM) molecules. To determine which ECM genes are expressed in the PM during LR patterning, we examined Zebrahub, a single-cell transcriptome atlas of developing zebrafish embryos^39^. We focused on the expression of ECM genes in the *tbx16*-positive PM cell cluster^40^, and found that several heparan sulfate proteoglycan (HSPG) core proteins including *gpc4*, *sdc2*, and *sdc4*, which are known to localize to the cell membrane^41^, were expressed in the PM at 12-15 ss (**Fig. 4a-c; Supplementary Fig. 8**). Enzymatic genes involved in synthesis and modification of HS sugar chains were also expressed in the PM cell cluster (**Supplementary Fig. 9a**). By contrast, the expression levels of chondroitin sulfate proteoglycan (CSPG) and dermatan sulfate proteoglycan (DSPG) core proteins were overall low (**Supplementary Fig. 9b**). ECM genes whose products mainly localize to the basement membrane, such as *agrn*, *fibronectin 1b (fn1b)*, *col5a2a*, *col11a1a*, and *col18a1a*, were also expressed in the PM (**Fig. 4b; Supplementary Fig. 9c**). Of them, HSPGs are known to form punctate clusters on the cell membrane and regulate the distribution and signalling range of secreted ligands^42–45^. Immunostaining with antibodies that recognize different types of HS sugar modifications^42^ revealed that HSPGs were abundantly distributed in the extracellular space of the zebrafish PM, colocalizing with Dand5-mV puncta (**Fig. 4d, e**).

**Figure 4.**
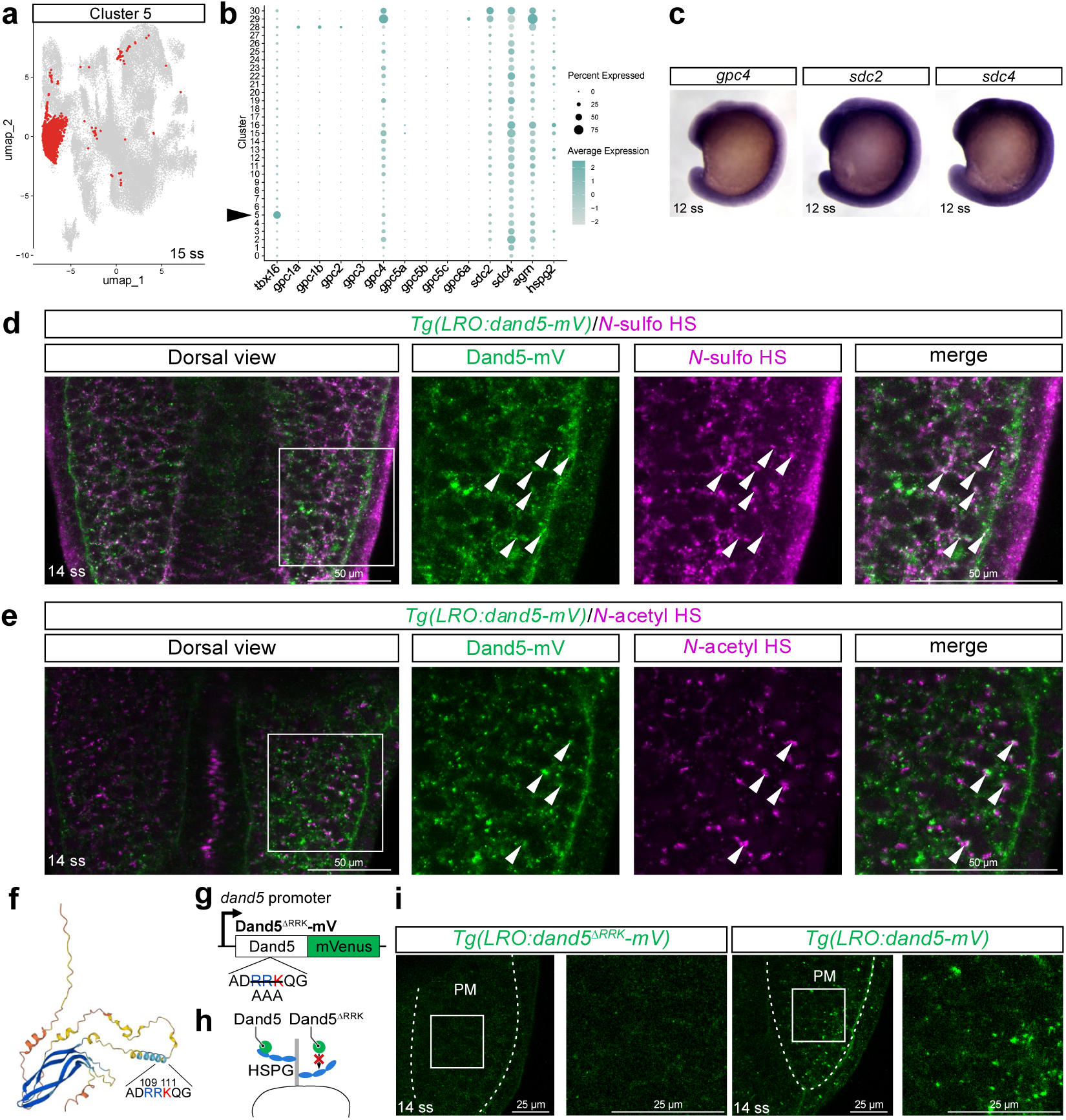
Involvement of HSPGs in the extracellular distribution of Dand5. **a,** UMAP analysis of cell clusters from 15-ss embryos. Datasets were retrieved from Zebrahub and subjected to reclustering. Cells belonging to cluster 5, which corresponds to the presomitic mesoderm (the posterior-most part of the PM) positive for *tbx16*, is highlighted in red. **b,** Dot plots representing the expression level of core proteins for HSPGs. Black arrowheads, PM cluster. **c,** Whole-mount *in situ* hybridization (WISH) of *gpc4, sdc2*, and *sdc4* in 12-ss embryos. **d, e,** Immunostaining of *N-*sulfo HS (d) and *N-*acetyl HS (e) in *Tg(LRO:dand5-mV)* embryos at 14 ss. Dorsal view of the tailbud. **f,** AlphaFold prediction of three-dimensional structure of zebrafish Dand5. **g,** Construction of Dand5^ΔRRK^-mV. The RRK motif is replaced with alanine. **h,** Schematic of the result in (i). **i,** Extracellular distribution of Dand5^ΔRRK^-mV in *Tg(LRO:dand5^ΔRRK^-mV)*. Dorsal view of the tailbud. White boxes, magnified areas. *n =* 6 (d), 6 (e), 9 (i) embryos.

To further test the interaction between Dand5 and HSPGs, we utilized the HSPG-degradation system in *Xenopus* embryos^29^, whose large cell size facilitates the visualization of subcellular localization of secreted proteins and HSPGs. We first confirmed that zebrafish Dand5-mV shows similar punctate distribution in *Xenopus* embryos (**Supplementary Fig. 10a, b**). To locally degrade HS chains near the Dand5-mV source, we expressed tethered-HepIII, a membrane-bound HS lyase that acts in a cell-autonomous manner^29^, by injecting its mRNA into adjacent blastomeres (**Supplementary Fig. 10c**). The Dand5-mV puncta were greatly diminished on the cells expressing tethered-HepIII (**Supplementary Fig. 10d**), suggesting that Dand5 interacts with HSPGs to accumulate on the cell surface.

We then determined the amino-acid motif in Dand5 that is essential for interaction with HSPGs. Because basic amino acid (arginine, histidine and lysine) residues are known to be important for electrostatic interaction with negatively-charged HS sugar chain^46–49^, we replaced motifs consisting of R (arginine) and K (lysine) in Dand5 with alanine (**Fig. 4f; Supplementary Fig. 11a**), and examined their distribution pattern using *Xenopus* embryos. The RRK (aa 109-111) mutant (Dand5^ΔRRK^-mV) showed decreased distribution, whereas the RERR (aa 214-217) mutant (Dand5^ΔRERR^-mV) displayed a punctuate distribution similar to that of the wild-type Dand5-mV (**Supplementary Fig. 11b**), suggesting that the RRK motif is involved in the interaction with HSPGs. We next established *Tg(LRO:dand5^ΔRRK^-mV)* zebrafish, in which the RRK motif in the transgene was replaced with alanine (**Fig. 4g**). Like in *Xenopus* embryos, the signal of Dand5^ΔRRK^-mV was lost in the PM of the *Tg(LRO:dand5^ΔRRK^-mV)* embryos, and was only detected in the LRO (**Fig. 4h, i; Supplementary Fig. 11c**). Dand5^ΔRRK^-mV accumulated in the PM upon morphotrap expression, suggesting that the mutation does not affect secretion (**Supplementary Fig. 11d**). These results show that the RRK motif is important for the interaction between Dand5 and HSPGs, and that punctate distribution of Dand5 in the PM is caused by interaction with HSPGs (**Fig. 4h**).

We then tested whether the ΔRRK mutation affects the activity of Dand5 as a Nodal inhibitor, by performing *in situ* hybridization for *spaw* in *Tg(LRO:dand5-mV)* and *Tg(LRO:dand5^ΔRRK^-mV)* embryos at 22 ss. We found that *spaw* expression was lost from the LPM of *Tg(LRO:dand5-mV)* embryos, in which Dand5-mV is symmetrically expressed in the LRO (**Supplementary Fig. 12**), confirming the activity of overexpressed Dand5-mV. Similarly, *spaw* expression was lost from the LPM of *Tg(LRO:dand5^ΔRRK^-mV)* embryos (**Supplementary Fig. 12**). These results demonstrate that Dand5^ΔRRK^ retains the inhibitory effect against Nodal.

### Diffusion of Spaw in the PM is essential for LR asymmetry formation in the LPM

Through reconstitutitve visualization, the Nodal-Gdf1 heterodimer was suggested to spread through the PM to the LPM, but the requirement of Nodal-Gdf1 diffusion for LR pattering in the LPM remained unclear. To test this, we engineered a novel trapping molecule called tethered-Dand5 by replacing the extracellular domain (anti-GFP VHH domain) of the morphotrap with Dand5 (**Fig. 5a**). Since Dand5 is known to physically interact with Nodal^14,20,50^, tethered-Dand5 was expected to trap endogenous Spaw, whereas the morphotrap only traps transgenic Spaw-mV (**Fig. 5a**). We first tested whether tethered-Dand5 can trap extracellular Spaw using *Xenopus* embryos (**Supplementary Fig. 13a**). We expressed Spaw-mV and tethered-Dand5 separately in *Xenopus* embryos by injecting their mRNAs into different blastomeres, and found that Spaw-mV secreted from their source cells was strongly accumulated on tethered-Dand5 expressing cells (**Supplementary Fig. 13b**). By contrast, tethered-Lefty1 expression in *Xenopus* embryos led to only weak accumulation of Spaw-mV (**Supplementary Fig. 13c**), showing the differential Spaw-trapping ability of Dand5 and Lefty. We then induced mosaic expression of tethered-Dand5 in the PM of the *Tg(LRO:spaw-mV)* embryos by injecting the *msgn1:tethered-dand5* plasmid (**Fig. 5b**). We found that Spaw-mV specifically accumulated on PM cells expressing tethered-Dand5 (**Fig. 5c, d**), suggesting that tethered-Dand5 traps Spaw proteins diffusing in the extracellular space of the PM. Furthermore, in wild-type embryos injected with the *msgn1:tethered-dand5* plasmid, up to 70% (59/84) of injected embryos lost *spaw* expression in the left LPM, while its expression in the LRO was maintained (**Fig. 5e, f**). This result indicates that the diffusion of Spaw (likely Spaw-Gdf1/3 heterodimer) through the extracellular space of the PM is essential for LR patterning in the LPM.

**Figure 5.**
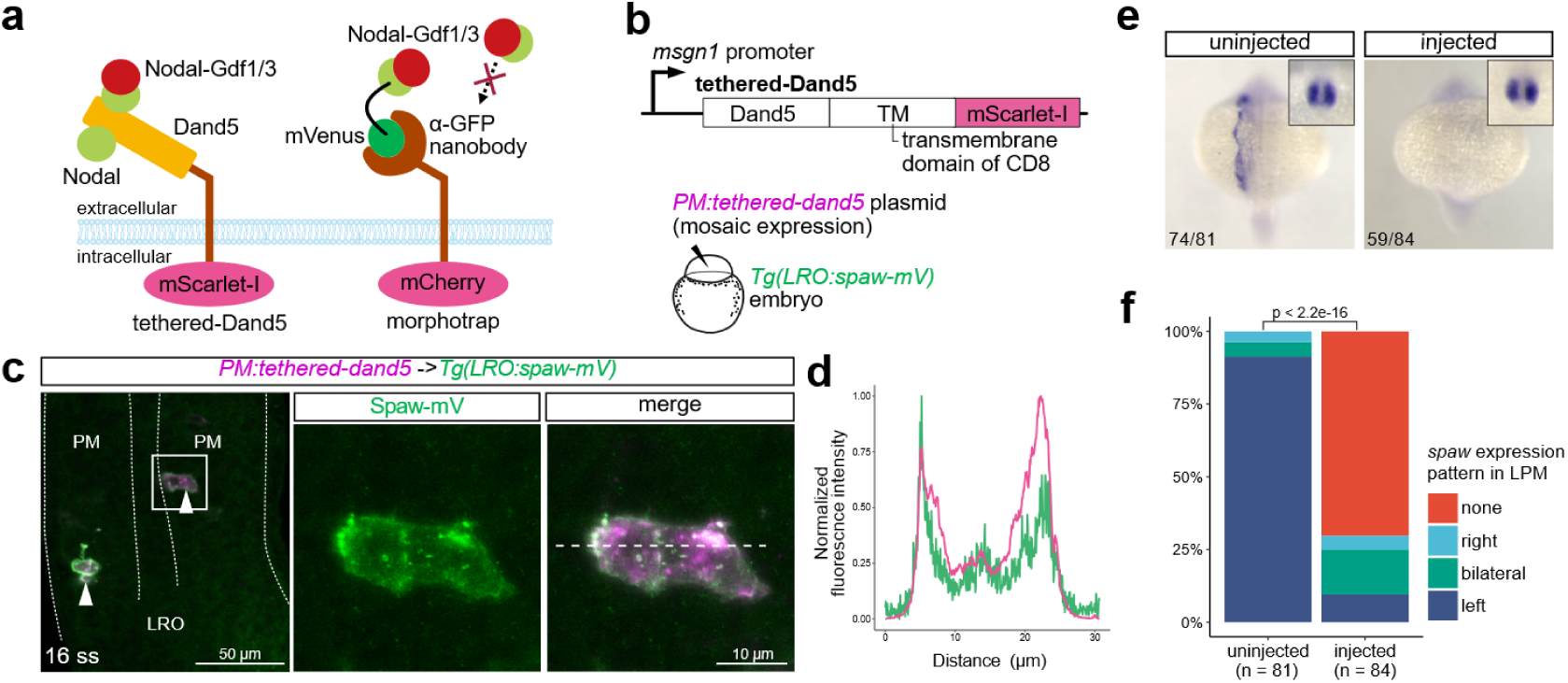
Trapping Spaw in the PM by tethered-Dand5. **a,** Schematic of tethered-Dand5 in comparison to the morphotrap. **b,** Construction of *PM:tethered-dand5* and experimental scheme in (c). **c,** Accumulation of Spaw-mV in the PM cells expressing tethered-Dand5 in a *Tg(LRO:spaw-mV)* embryo injected with the *PM:tethered-dand5* plasmid. Dorsal view of the tailbud. White arrowheads, tethered-Dand5-positive cells. *n* = 10 embryos. **d,** Normalized intensity plot of Spaw-mV and tethered-Dand5 fluorescence. Fluorescence intensity was measured along the white dotted line in (d). **e, f,** Increased frequency of LR defects in *PM:tethered-dand5*-injected embryos. Wildtype embryos were injected with the *PM:tethered-dand5* plasmid at 1-cell stage, harvested at 22-ss stage, and subjected to WISH for *spaw* (e). Embryos were classified according to the expression pattern of *spaw* in the LPM (f). Insets in (e), expression of *spaw* in the LRO. *p*-value in (f) was calculated using Fisher’s exact test.

## Discussion

Visualization and manipulation of secreted ligands in the extracellular space is essential to understand their modes of action in various developmental contexts. Our reconstitutive imaging system together with ligand-trapping technologies revealed the distinct behaviors of secreted ligands and inhibitors related to LR asymmetry formation, Nodal, Gdf1, Dand5 and Lefty, in zebrafish embryos (**Fig. 6**). Based on our data, we discuss how these LR-related ligands and inhibitors act to induce asymmetric gene expression in the LPM.

**Figure 6.**
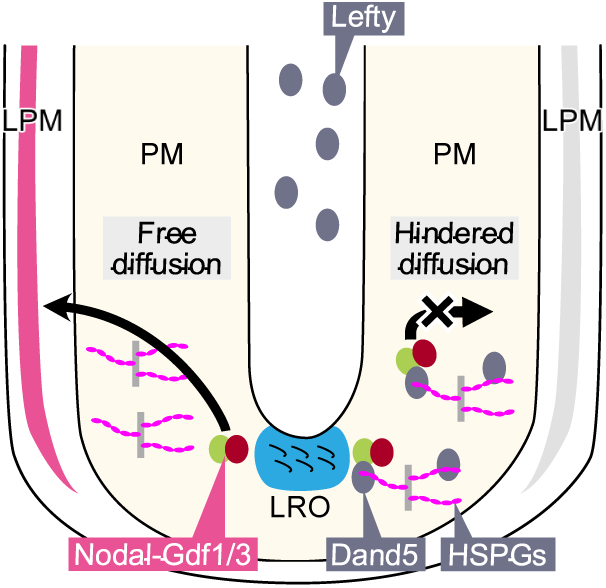
Proposed model of extracellular roles of Nodal-Gdf1, Dand5 and Lefty during LR asymmetry formation in the zebrafish embryo. The Nodal-Gdf1 heterodimer is secreted from the LRO and spreads to the LPM through the PM. Dand5 is secreted from the right side of the LRO and distributes asymmetrically on the right side of the PM through interaction with HSPGs.

### Extracellular behavior of Nodal-Gdf1 during LR asymmetry formation

Our visualization and manipulation experiments demonstrated that the Nodal-Gdf1 heterodimer produced in the LRO exerts its long-range effect by diffusing through the PM (**Figs. 2**, **5**). Extracellular distribution of Nodal and Gdf1 during mesoderm induction has been examined by mRNA injection experiments^8,51^. At the blastula stage (i.e., mesoderm induction period) of zebrafish embryos, overexpressed Nodal is uniformly distributed in the extracellular space of the blastoderm^8,9^. By contrast, in the present study, the extracellular distribution of Spaw-mV, which was expressed in the LRO under the control of the *dand5* promoter, was detected only when they were trapped by the morphotrap or tethered-Dand5 (**Figs. 1**, **5**). This was also the case with sec-mV (**Fig. 1**), which is known to have low affinity for the ECM in *Xenopus* embryos^29^. Thus, Spaw-mV (and Spaw-mV-Gdf1/3-V5; **Fig. 2**) appears to be weakly associated with ECM molecules in the PM, and instead spread in a free-diffusion-like manner at concentrations below the detection threshold. Consistent with our observation, Oki et al. (2007) overexpressed Myc-tagged Nodal in the mouse node (LRO), but its extracellular signal was hardly detected. The free-diffusion nature of Nodal ligands is thus conserved between mouse and zebrafish, in sharp contrast with other signalling ligands such as Wnt, FGF and Hedgehog, which have strong affinity for ECM molecules including HSPGs^42,52,53^.

Our results highlight the importance of the extracellular space in the PM as a diffusion path for Nodal-Gdf1. Previous studies in mouse and *Xenopus* embryos suggested that intercellular calcium signalling through gap junctions in the endoderm is involved in signal transmission from the LRO to the LPM^54–56^. However, our trapping experiments by the morphotrap and tethered-Dand5 (**Figs. 2**, **5**) indicate that Nodal travels through the PM to the LPM as a heterodimer with Gdf1, thereby acting as a major signal transducer from the LRO, at least in zebrafish embryos. Despite the presence of Nodal and its receptors, ActRIIB and ALK4 (**Supplementary Fig. 14**), the PM remains symmetric throughout development. Notably, the expression of Oep, a Nodal coreceptor essential for Nodal signalling^57^, is restricted to the notochord and LPM (**Supplementary Fig. 14**). In *MZoep* mutants, the distribution of Nodal was expanded in the blastula embryo^58^, possibly due to reduced internalization of Nodal ligands. During LR asymmetry formation, Nodal-Gdf1 may similarly pass through the PM owing to the low expression level of Oep, thereby specifically targeting the LPM.

### Extracellular behaviors of Dand5 and Lefty during LR asymmetry formation

Since *nodal* is bilaterally expressed around the LRO, its activity must be repressed on the right side of the embryo to generate LR asymmetry in the LPM. Like *nodal*, *dand5* is initially expressed bilaterally in the LRO, but the leftward flow generated by the LRO cilia induces 3’-UTR-dependent degradation of *dand5* mRNA on the left side^37,38,59^. In addition to such asymmetry at the mRNA level, we demonstrated for the first time that asymmetric *dand5* mRNA expression in the LRO leads to asymmetric extracellular distribution of Dand5 protein in the PM (**Fig. 3**). Dand5 is probably secreted from the basal side of LRO (KV)-epithelial cells, which lack the basement membrane^26^, and spreads directly to the adjacent PM (**Fig. 3; Supplementary Fig. 7**). Asymmetric *dand5* expression in the LRO is, therefore, directly reflected by the asymmetric distribution of Dand5 protein in the PM. This may represent a conserved step of LR asymmetry formation across vertebrates, because asymmetric *dand5* expression in the LRO has been observed from teleosts to mammals^20,25,37^.

The characteristic distribution pattern of Dand5 in the PM (**Fig. 3**) is attributed to its affinity for HSPGs, ECM components that are distributed in a punctate manner on the cell surface^42^ (**Fig. 4**). Involvement of HSPGs in LR asymmetry formation has been shown in *Xenopus* embryos, where HSPGs degradation at the gastrula stage leads to the randomization of LR axis^60^. Our results suggest that HSPGs participate in LR asymmetry formation in zebrafish embryos at early segmentation stages by interacting with Dand5. How, then, do HSPGs modulate the mode of action of Dand5? We showed that the interaction with HSPGs is not essential for Dand5 to function as a Nodal inhibitor, as Dand5 lacking the HS-interacting motif retained its inhibitory effect against Nodal (**Supplementary Fig. 12**). One possibility is that HSPGs restrict the distribution of Dand5 to the right PM, allowing them to act as a local and asymmetric Nodal inhibitor. Additionally, HSPGs may promote the internalization and turnover of the Nodal-Dand5 complex, as cell-surface HSPGs such as Syndecans and Glypicans are frequently internalized^42^.

The extracellular behavior of Dand5 contrasts with that of Lefty, which is a fast-diffusing molecule that hardly interacts with Nodal ligands and the ECM^6,61^ (**Figs. 3**, **5**). Lefty is secreted from the midline tissues and inhibits Nodal signal in the right LPM by competitively binding to Nodal receptors^61^. Unlike *lefty*, *dand5* expression diminishes by the time when *nodal* starts to be expressed in the left LPM^26^. Considering those characteristics, Lefty likely acts as a long-range suppressor of Nodal ligands in the right LPM, whereas Dand5 serves as a local suppressor that primarily targets Nodal ligands from the LRO.

In conclusion, our results reveal a unique mode of action for LR-related ligands and inhibitors in the extracellular space of the PM in zebrafish embryos. Nodal-Gdf1 freely spreads on the left side, whereas Dand5 accumulates in the ECM and locally inhibits Nodal signalling on the right side. This asymmetric interaction of ligands and inhibitors in the extracellular space of the PM provides a new framework for understanding the mechanisms underlying LR asymmetry formation in vertebrate embryos.

## Materials and Methods

### Zebrafish strain and manipulation of embryos

The RW (RIKEN WT) and TL2E (Tüpfel long fin 2E) strains were used as the wild-type zebrafish strains. Adult fish and embryos were maintained under standard conditions. Embryos were incubated at 23-28 °C in 1/3× Ringer’s solution (38.7 mM NaCl, 0.97 mM KCl, 1.67 mM HEPES, 1.80 mM CaCl_2_) until they reach the stage of interest. Staging of embryos is based on Kimmel et al. (1995). All experimental procedures and animal care were carried out according to the animal ethics committee of the University of Tokyo (Approval No. 20-2) and Kyoto Sangyo University (Approval No. 2025-25).

### Manipulation of *Xenopus* embryos

Manipulation of *Xenopus laevis* embryos and microinjection experiments were carried out according to standard methods^63,64^. Briefly, unfertilized eggs were obtained from female frogs injected with gonadotropin (ASKA Pharmaceutical), and artificially fertilized using testis homogenate. Fertilized eggs were dejellied with 2% L-cysteine solution (pH 7.8) and incubated in 0.1×Steinberg’s solution at 14-18 °C. Embryos were staged according to Nieuwkoop and Faber (1967)^65^.

### Plasmid construction

Coding sequences (CDSs) of Spaw, Gdf1/3, Dand5, and Lefty1 were cloned from zebrafish one-cell and 16-ss cDNAs, and was inserted to pCSf107mT^63^. mVenus CDS was inserted two amino acids downstream of the putative prodomain cleavage site (RRHKR for Spaw and RRKR for Gdf1/3) to construct pCSf107-Spaw-mV and -Gdf1/3-mV. pCSf107-Dand5-mV was constructed by fusing mVenus to the C-terminus of Dand5 with a 5×Gly linker. Then, the 5-kb *dand5* promoter^26^ and Spaw-mV, Gdf1/3-mV or Dand5-mV were assembled into the pDestTol2pA2 vector to construct pDestTol2pA2*-dand5:spaw-mV*, -*dand5:gdf1/3-mV*, and -*dand5:dand5-mV*, respectively. The signal peptide of Spaw (aa 1-19) was fused to the N-terminus of mVenus to construct pDestTol2pA2-*dand5*:sec-mVenus. 4× HA tags were inserted after the signal peptide sequence of Spaw-mV in pDestTol2pA2*-dand5:spaw-mV* to construct pDestTol2pA2*-dand5:4HA-spaw-mV*. pDestTol2pA2*-drl:morphotrap*, pBSSK-*msgn1:morphotrap*, and pDestTol2pA2*-dand5:morphotrap* were constructed by replacing the reporter CDSs of pDestTol2pA2*-drl:EGFP*^30^, pBSSK-*msgn1:mCherry*^32^, and pDestTol2pA2*-dand5:EGFP*^26^, respectively, with the morphotrap CDS amplified from pCSf107-morphotrap^29^. To construct pBSSK-*dand5:spaw-V5-αcry:GFP* and pBSSK-*dand5:gdf1/3-V5-αcry:GFP*, pCSf107-Spaw-V5 and pCSf107-Gdf1/3-V5 were constructed by inserting a V5 epitope tag at two amino acids downstream of their cleavage sites. Then, Spaw-V5 or Gdf1/3-V5 sequence was assembled with the *dand5* promoter and the backbone sequence of pBSSK-*msgn1:CreERT2*-*αcry:GFP*^66^. The 547-bp 3’UTR sequence of *dand5* was inserted into pDestTol2pA2-*dand5*:*dand5-mV* to construct pDestTol2pA2-*dand5:dand5-mV*-3’UTR. Dand5-OLLAS and Dand5-mEosEM were constructed by fusing the OLLAS epitope tag^67^ and mEosEM^36^, respectively, to the C-terminus of Dand5. Alanine mutants of Dand5-mV (Dand5^ΔRRK^-mV, Dand5^ΔRERR^-mV, and Dand5^ΔRRKΔRERR^-mV) were constructed by PCR-based mutagenesis. Tethered-Dand5 and tethered-Lefty1 were constructed by assembling Dand5 or Lefty1, the transmembrane domain of mouse CD8^5^, and mScarlet-I^68^. pBSSK(+)-tol2-*msgn1:tethered-Dand5* and pBSSK(+)-tol2-*msgn1:tethered-Lefty1* were constructed by replacing mCherry CDS in pBSSK-*msgn1:mCherry* with tethered-Dand5 or tethered-Lefty, respectively. Cloning and subcloning was performed using PrimeSTAR GXL polymerase (Takara Bio), PrimeSTAR MAX polymerase (Takara Bio), In-Fusion Snap Assembly kit (Takara Bio) and NEBuilder HiFi DNA Assembly kit (New England Biolabs). Sequences of primers used for cloning are listed in Supplementary Table. Digital plasmid maps are available upon request.

### Transgenesis

Zebrafish transgenesis was carried out by the Tol2 transposase method^69^. To obtain founders, Tol2 mRNA (25 ng/µL) and plasmids (12.5 ng/µL) were injected into one-cell stage embryos of the RW strain. Founders carrying transgenes were screened by crossing with the TL2E strain.

### Immunostaining

Embryos were fixed with 4% paraformaldehyde (PFA) in PBST at 4 °C for overnight or at 25 °C for two hours, washed with PBST for three times and were stored at 4 °C until use. Fixed embryos were permeabilized with 1% Triton X-100 in PBS, blocked with 2% BSA in PBSDT (PBS/1% DMSO/0.1% Triton X-100) and were immunostained with following antibodies: mouse monoclonal anti-GFP (1/50, A-11120, Invitrogen), rabbit polyclonal anti-GFP (1/800, 632592, Clontech), mouse monoclonal anti-V5 (1/500, V8012, Sigma-Aldrich), rat monoclonal anti-HA (1/400, ROAHAHA, Roche), and rat monoclonal anti-OLLAS (1/500, MA5-16125, Invitrogen), anti-*N-*acetyl HS (1/50, NAH46, Seikagaku), anti-*N-*sulfo HS (1/200, HepSS-1, Seikagaku). Secondary antibodies for immunofluorescence were anti-rabbit IgG labelled with Alexa Fluor 488 or 568 (A-11008 or A-10042, Invitrogen), anti-mouse IgG labelled with Alexa Fluor 488, 568 or 647 (A-11001, A-10037 or A-21236, Invitrogen), anti-rat IgG labelled with Alexa Fluor 405 (ab175670, abcam), and anti-mouse IgM labelled with Alexa Fluor 555 (A-21426, Invitrogen). For counterstaining cell membranes, Phalloidin labelled with Alexa Fluor plus 647 (A30107, Invitrogen) was added to the secondary antibody solution. Stained embryos were mounted in 1% LMP agarose/PBS (16520-050, Invitrogen) in a glass-based dish (3911-035, Iwaki) and imaged using LSM710 and LSM 980 confocal microscopes (Carl Zeiss). 40× and 25× water-immersion objective lenses (LD C-Apochromat 40x/1.1 W Korr M27 and LD LCI Plan-Apochromat 25x/0.8 Imm Corr DIC M27, Carl Zeiss) were used. Contrast adjustment, cropping of captured images, and intensity quantification were performed using Fiji/ImageJ ^70^ (NIH). For counting Dand5-mV puncta (**Fig. 3h**), background signals were first subtracted from the raw confocal images using the *Subtract Background* command. The images were then converted to binary using the *Threshold* command, followed by segmentation with the watershed algorithm. The number of particles within a 76.80 × 100.46 µm area in the left and right PM was quantified using the *Analyze Particles* command.

### *in situ* hybridization

Whole-mount *in situ* hybridization (WISH) was carried out according to Yamaguchi et al. (2018)^71^ with slight modifications. Hybridization was performed overnight at 60 °C and BM Purple (Roche) was used for chromogenic reaction. Digoxigenin (DIG)-labelled RNA antisense probes were synthesized from template plasmids, pCSf107-*sdc2*-T, pCSf107-*sdc4*-T, and pCSf107-*gpc4*-T, which contain full-length CDS of each gene, using T7 RNA polymerase and DIG RNA Labeling Mix (Roche). Fluorescence *in situ* hybridization was performed according to Ikeda et al. (2022)^26^. Chromogenic reaction was performed using the TSA Plus Cyanine 3 Kit (Akoya Biosciences).

### Alphafold

Three-dimensional structure of zebrafish Dand5 (UniProtKB accession number: Q76C29) was predicted by Alphafold^72,73^. Snapshots of predicted structure (AFDB accession number: AF-Q76C29-F1-v4) were retrieved from AlphaFold Protein Structure Database (https://alphafold.ebi.ac.uk/), and processed by Affinity Publisher (Serif Europe).

### Zebrahub

Filtered count matrices for single-cell RNA-sequencing data from four 15-ss zebrafish embryos (TDR39-42) were retrieved from the Zebrahub portal site^39^ (https://zebrahub.sf.czbiohub.org/). *Seurat* package (version 5.2.1) in *R* were used for data processing and visualization^74,75^. Cells expressing less than 200 or more than 9,000 genes were removed from the dataset. Dimension reduction was conducted by Principal Component Analysis (PCA) based on 2,000 highly variable genes, which was determined by *FindVariableFeatures* function. Four replicates were then integrated by Canonical Correlation Analysis (CCA), using *IntegrateLayers* function with the argument “method = CCAIntegration.” Clustering of cell types was performed by *FindNeighbors* and *FindClusters* function using the first 20 components in CCA integration. Visualization of cell clusters were performed by UMAP algorithm^76^. Dot plots representing the expression level and frequency of genes of interest in each cluster were drawn by *DotPlot* function.

## Author Contributions

Conceptualization, T.I. and H.T.; Methodology, T.I., T.K., Y.M., T.Y., J.P. and R.B.; Investigation, T.I.; Resources, Y.M. and T.Y.; Writing - Original Draft, T.I.; Writing - Review & Editing, all authors; Supervision, H.T.; Funding Acquisition, T.I., T.K. and H.T.

### Acknowledgement

We thank Dr. Christian Mosimann for *drl:EGFP* (pCM298) plasmid; Seikagaku corporation for NAH46 antibody; Dr. Osamu Yoshie for HepSS1 antibody; Dr. Masanori Taira for providing *Xenopus* facility and discussion; Dr. Shinji Takada for providing zebrafish facility and discussion; Yuki Yamagishi and Ikuko Fukuda for fish husbandry. This research was supported by Grant-in-Aid for Japan Society for the Promotion of Science (JSPS) Fellows under Grant Numbers JP18J21960 (T.I.), JSPS KAKENHI under Grant Numbers JP22K20625 (T.I.), JP23K14190 (T.I.), and JP24H01982 to H.T., and Joint Research of the Exploratory Research Center on Life and Living Systems (ExCELLS) under ExCELLS program numbers 24EXC340, 23EXC322, 22EXC311, and 20-320).

## Supplementary Figure Legends

**Supplementary Figure 1. Accumulation of Spaw-mV in the morphotrap-expressing LPM**

Confocal images of *Tg(LRO:spaw-mV);Tg(LPM:morphotrap)* embryos at 12 ss, 14 ss and 16 ss. Maximum intensity projections of each image are shown. Dorsal views of the tailbud. White arrowheads, signal of Spaw-mV in the morphotrap-expressing LPM. *n =* 3 to 5 embryos per stage.

**Supplementary Figure 2. Detection of Spaw prodomain**

**a,** Construction of *LRO:HA-spaw-mV*. 4× HA tags are inserted between the signal peptide (SP) and the prodomain of Spaw-mV. **b,** Immunostaining of a 20-ss *LRO:HA-spaw-mV;LPM:morphotrap* embryo by anti-GFP and anti-HA antibodies. Lateral view of the tailbud. Maximum intensity projection is shown. White dotted lines, contour of embryo. White boxes, magnified areas. *n =* 3 embryos. **c,** Fluorescence intensity of HA-Spaw-mV and morphotrap along the anterior-posterior axis from the LRO to the LPM (yellow line in b). Green, mVenus. Magenta, morphotrap. Black, HA. Intensity was normalized so that the maximum value is 1 and the minimum value is 0 for each channel. **d,** Schematic of the result in (b).

**Supplementary Figure 3. Accumulation of Spaw-mV in the morphotrap-expressing PM**

*Tg(LRO:spaw-mV);Tg(PM:morphotrap)* 20-ss embryo. White dotted lines, contour of embryo. Lateral view of the tailbud. Maximum intensity projection is shown. White boxes, magnified areas. PSM, presomitic mesoderm. sm, somite. *n =* 6 embryos.

**Supplementary Figure 4. Morphotrap expression in the hatching gland of *Tg(LRO:spaw-mV)* embryo**

Morphotrap expression in the hatching gland of *Tg(LRO:spaw-mV)* embryo. *Tg(LRO:spaw-mV)* embryos were injected with *he1.1:morphotrap* plasmid at 1-cell stage, and were imaged at 20 ss. Maximum intensity projection is shown. No Spaw-mV signal was detected on the hatching-gland cells expressing the morphotrap. White dotted lines, contour of embryo. White boxes, magnified areas. PSM, presomitic mesoderm. HG, hatching gland. nc, notochord. *n =* 5 embryos.

**Supplementary Figure 5. in situ sandwich assay in Tg(LRO:spaw-mV); Tg(LRO:spaw-V5);Tg(LPM:morphotrap)**

**a, b,** 22-ss embryos of *Tg(LRO:spaw-mV);Tg(LRO:spaw-V5);Tg(LPM:morphotrap)* (a) and *Tg(LRO:spaw-V5);Tg(LPM:morphotrap)* (b) immunostained with anti-GFP and anti-V5 antibodies. Lateral view of the tailbud. Maximum intensity projections of each image are shown. White dotted lines, contour of embryo. White boxes, magnified areas. **c, d,** Fluorescence intensity of Spaw-mV, Spaw-V5 and morphotrap along the yellow lines in (a) and (b) was plotted in (c) and (d), respectively. Green, Spaw-mV. Red, Spaw-V5. Magenta, morphotrap. *n =* 3 embryos for each.

**Supplementary Figure 6. Distribution of Dand5-mV and HSPGs at 20 ss**

20-ss *Tg(LRO:dand5-mV)* embryos immunostained with anti-*N-*sulfo HS (A) or anti-*N-*acetyl HS (B) antibodies. *N-*acetyl HS strongly accumulated around the notochord, whereas *N*-sulfo HS was poorly detected. Lateral view of the tailbud. White boxes, magnified areas. Black arrowheads, accumulation of Dand5-mV around the notochord (nc). *n =* 3 embryos each.

**Supplementary Figure 7. Extracellular distribution of Dand5-OLLAS and Dand5-mEosEM**

**a,** *Tg(LRO:dand5-OLLAS)*. The OLLAS epitope tag is fused to the C-terminus of Dand5. Dorsal view of the tailbud. The embryo was counterstained with phalloidin to highlight cell membranes. White boxes, magnified area. White arrowheads, Dand5-OLLAS puncta on the cell surface. Black arrowheads, Dadn5-OLLAS puncta in the cytoplasm. *n =* 8 embryos. **b,** *Tg(LRO:dand5-mEosEM)*. mEosEM is fused to Dand5 in its C-terminus. Signals in the KV lumen and PM are shown. Note that Dand5-mEosEM is abundant in the KV fluid, suggesting that Dand5 is secreted from the apical side of KV-epithelial cells into the lumen, as well as from the basal side into the PM. White arrowheads, punctate localization of Dand5-mEosEM in the PM.

**Supplementary Figure 8. scRNA-seq data for *gpc4*, *sdc2* and *sdc4***

**a,** UMAP plots representing cell cluster of 15-ss zebrafish embryos. Black arrowheads indicate cluster 5, which corresponds to the presomitic mesoderm (**Fig. 4a**). **b-e,** UMAP plots representing the expression level of the premositic-mesoderm marker *tbx16* (b), *gpc4* (c), *sdc2* (d), and *sdc4* (e).

**Supplementary Figure 9. scRNA-seq data for CSPGs/DSPGs core proteins, Fibronectin, Laminin, and HSPGs-related enzyme genes**

**a-c,** Dot plots representing the expression level of genes encoding CSPGs/DSPGs core proteins (a), Fibronectin and Laminin (b), and HSPGs-related enzymes (c). *tbx16* is a marker for the presomitic mesoderm.

**Supplementary Figure 10. Interaction of Dand5 and HSPGs in *Xenopus* embryos**

**a,** Experimental scheme for (b). mRNAs encoding *dand5-mV* (500 pg) were injected into a dorsal blastomere of the 4-cell stage *Xenopus* embryos. Confocal images were taken at the gastrula (st. 10-10.5). **b,** Distribution of Dand5-mV in a *Xenopus* embryo. Embryos were injected with 500 pg of *dand5-mV* mRNA. Animal region. **c,** Experimental scheme for (d). mRNAs encoding *dand5-mV* (500 pg) or *tethered-HepIII* (100 pg) were injected into two different dorsal blastomere of the 4-cell stage *Xenopus* embryos. *mECFP* mRNA (400 pg) was co-injected with *tethered-HepIII* mRNA as a lineage tracer. Confocal images were taken at the gastrula (st. 10-10.5). **d,** Decrease of Dand5-mV distribution by tethered-HepIII expression. White arrowheads, tethered-HepIII expressing cells. Black arrowheads, tethered-HepIII non-expressing cells. White asterisks, source cells of Dand5-mV. *n =* 8 (b) and 4 (d) embryos.

**Supplementary Figure 11. HS-binding motif in Dand5**

**a,** Amino acid sequence of zebrafish Dand5. Arginine (R) and lysine (K) residues are marked in blue and red, respectively. Two motifs rich in R and K underlined in green were replaced with alanine. **b,** Extracellular distribution analysis of Dand5 alanine mutants in *Xenopus* embryos. mRNAs encoding *dand5-mV*, *dand5^ΔRRK^-mV* or *dand5^ΔRERR^-mV* (500 pg each) were injected into a dorsal blastomere of the 4-cell stage *Xenopus* embryos. Confocal images were taken at the gastrula (st. 10-10.5). *n =* 7 (Dand5-mV), 9 (Dand5^ΔRRK^-mV), 11 (Dand5^ΔRERR^-mV) embryos. White asterisks, source cells. White arrowheads, extracellular signal of Dand5-mV. **c,** Expression of Dand5^ΔRRK^-mV in KV-cells of the 14-ss *Tg(LRO:dand5^ΔRRK^-mV)* embryo. Embryo is counterstained with phalloidin to highlight cell membranes. Dorsal view. *n =* 3 embryos. **d,** Accumulation of Dand5^ΔRRK^-mV in the PM of the 16-ss *Tg(LRO:dand5^ΔRRK^-mV);Tg(PM:morphotrap)* embryo. Dorsal view. *n =* 14 embryos.

**Supplementary Figure 12. LR defects in *Tg(LRO:sec-mV)*, *Tg(LRO:dand5-mV)* and *Tg(LRO:dand5^ΔRRK^-mV)* embryos**

22-ss embryos of *Tg(LRO:sec-mV)*, *Tg(LRO:dand5-mV)* and *Tg(LRO:dand5^ΔRRK^-mV)* were subjected to WISH for *spaw* to test the effects of each transgene on LR patterning in the LPM. As in wildtype embryos, *spaw* was expressed in a left-side specific manner in *Tg(LRO:sec-mV)* embryos. By contrast, *spaw* expression was not detected in *Tg(LRO:dand5-mV)* and *Tg(LRO:dand5^ΔRRK^-mV)* embryos, demonstrating the suppressive effects of Dand5-mV and Dand5^ΔRRK^-mV on Nodal signalling.

**Supplementary Figure 13. Tethered-Dand5 and Tethered-Lefty1 expression in *Xenopus* embryos**

Spaw trapping experiment in *Xenopus* embryos. *spaw-mV* mRNA and *tethered-dand5* (b) or *tethered-dand5* (c) mRNA (500pg each) were injected into two different dorsal blastomeres of the 4-cell stage embryos (a). Black arrowheads in (b), Accumulation of Spaw-mV on tethered-Dand5 expressing cells. White arrowheads in (c), Less accumulation of Spaw-mV on tethered-Lefty1 expressing cells. Asterisk, Spaw-mV source cells. *n =* 8 embryos for (b) and (c).

**Supplementary Figure 14. Expression of Nodal receptors in the PM**

**a-c,** Fluorescence *in situ* hybridization of the Nodal coreceptor *oep* (a), Nodal receptors *acvr1ba (alk4)* (b), and *acvr2ba (actr2b)* (c) in 12-ss embryos. Dorsal view of the tailbud. Maximum intensity projections of each image are shown. White dotted lines, contour of embryo. nc, notochord. *n =* 3 embryos per gene.

