## Supplementary figures for "Differential spreading behaviors of Nodal signalling molecules in the extracellular space cooperatively shape left-right asymmetry"

### Supplementary Figure 1

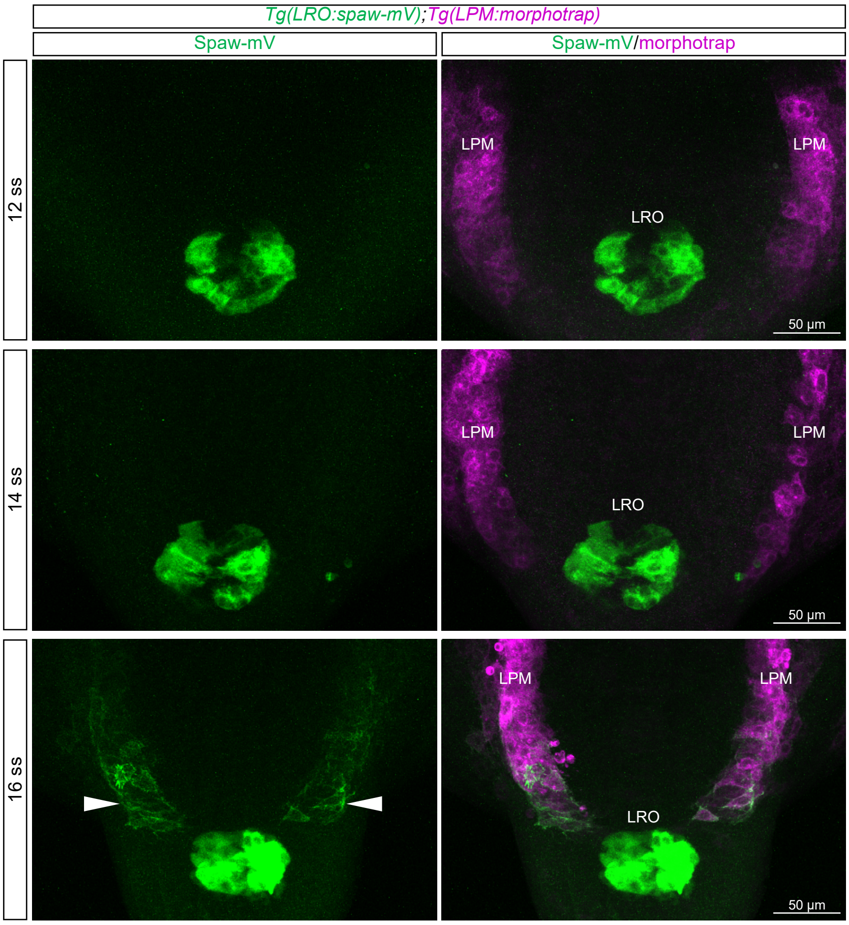

### Supplementary Figure 2

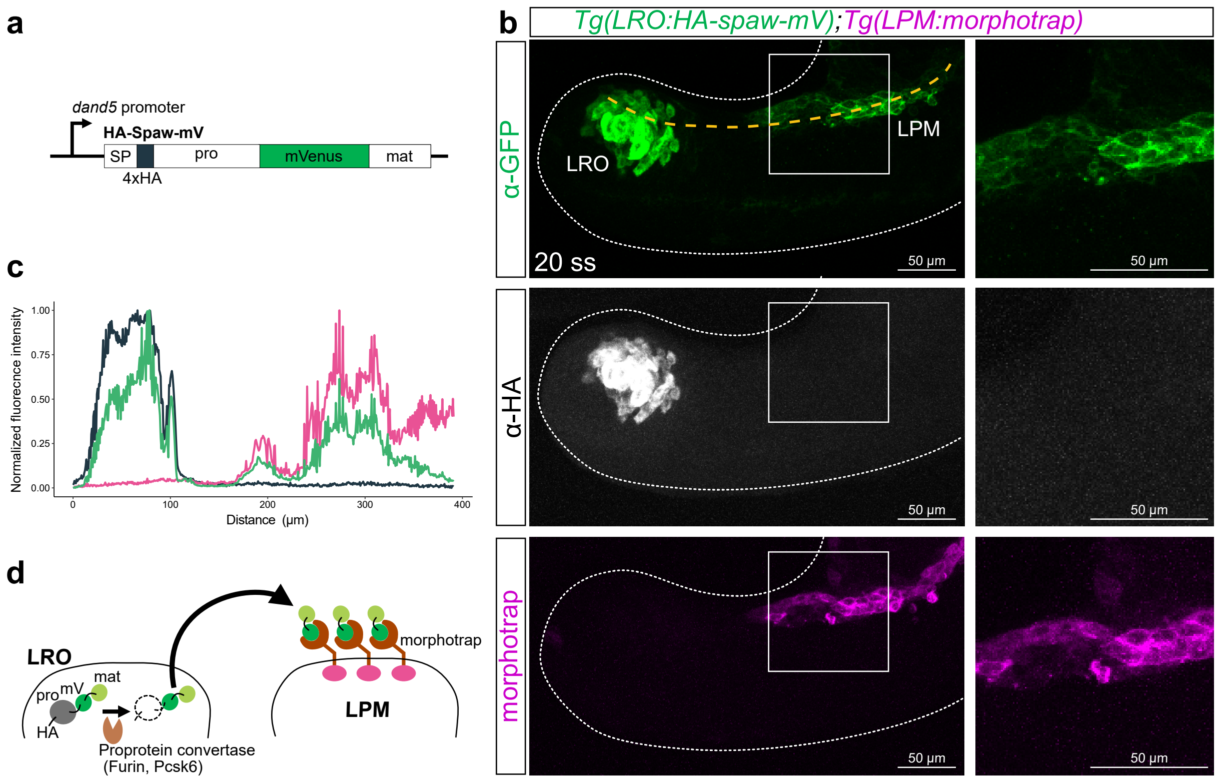

### Supplementary Figure 3

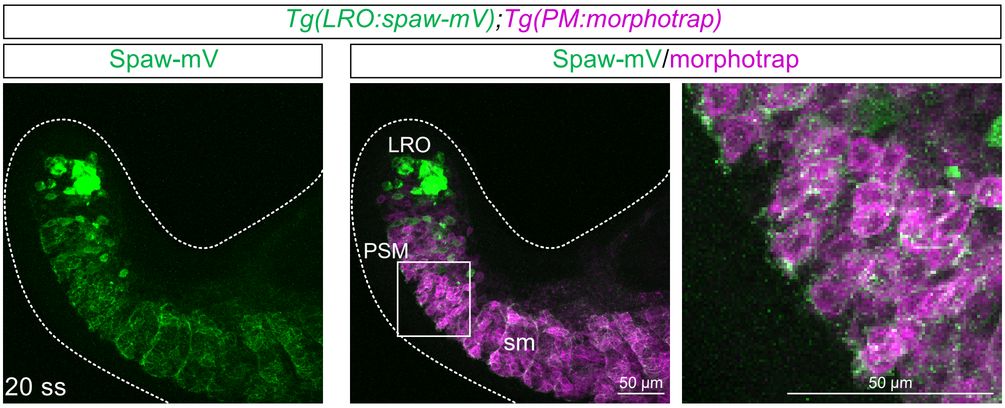

### Supplementary Figure 4

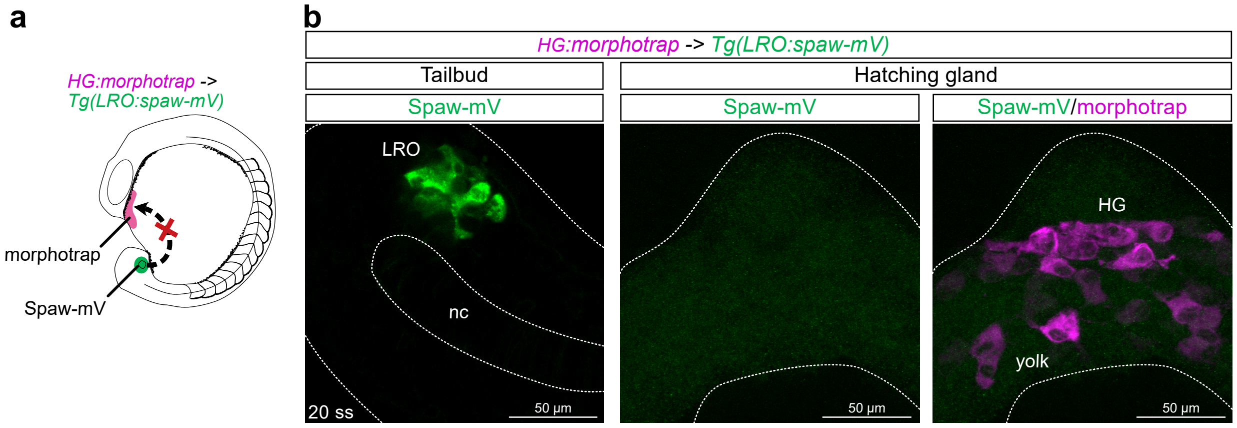

### Supplementary Figure 5

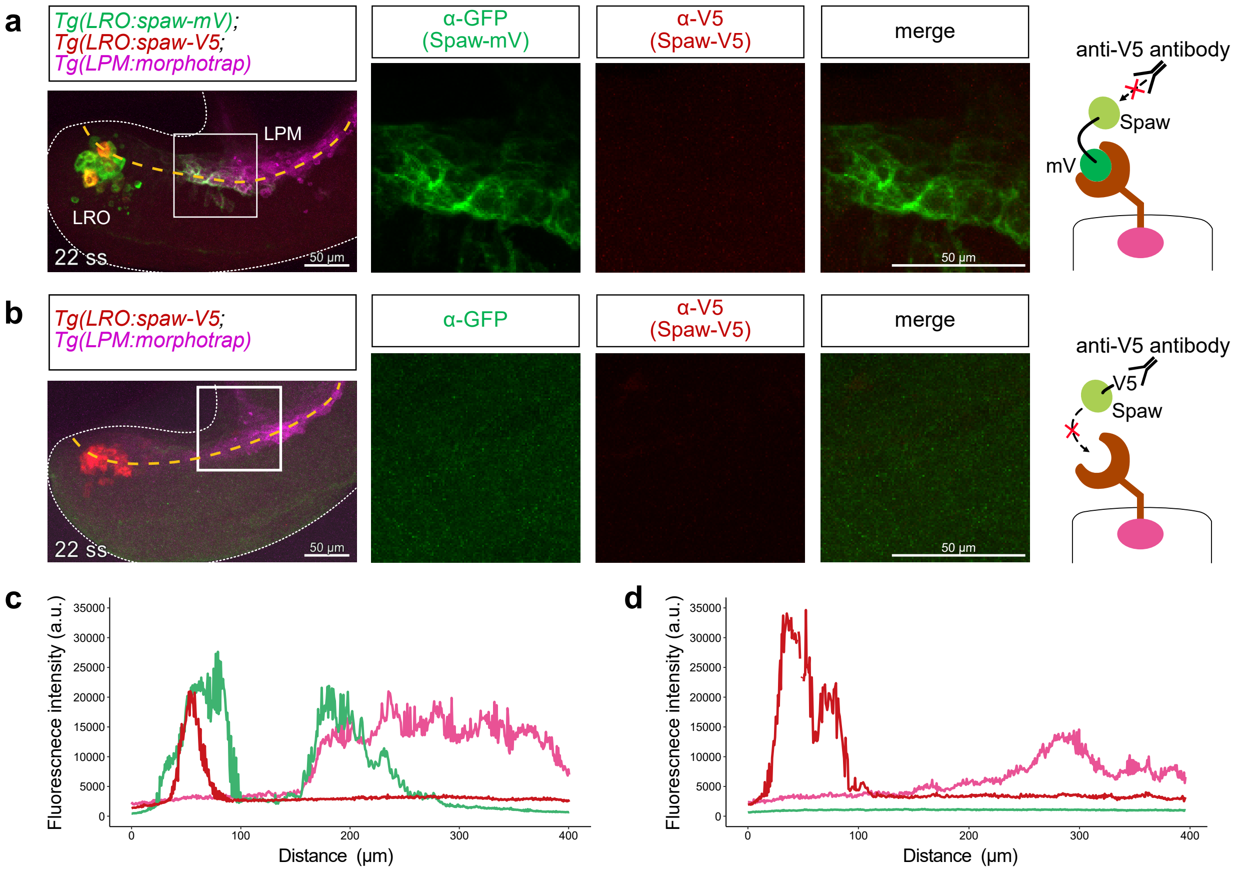

### Supplementary Figure 6

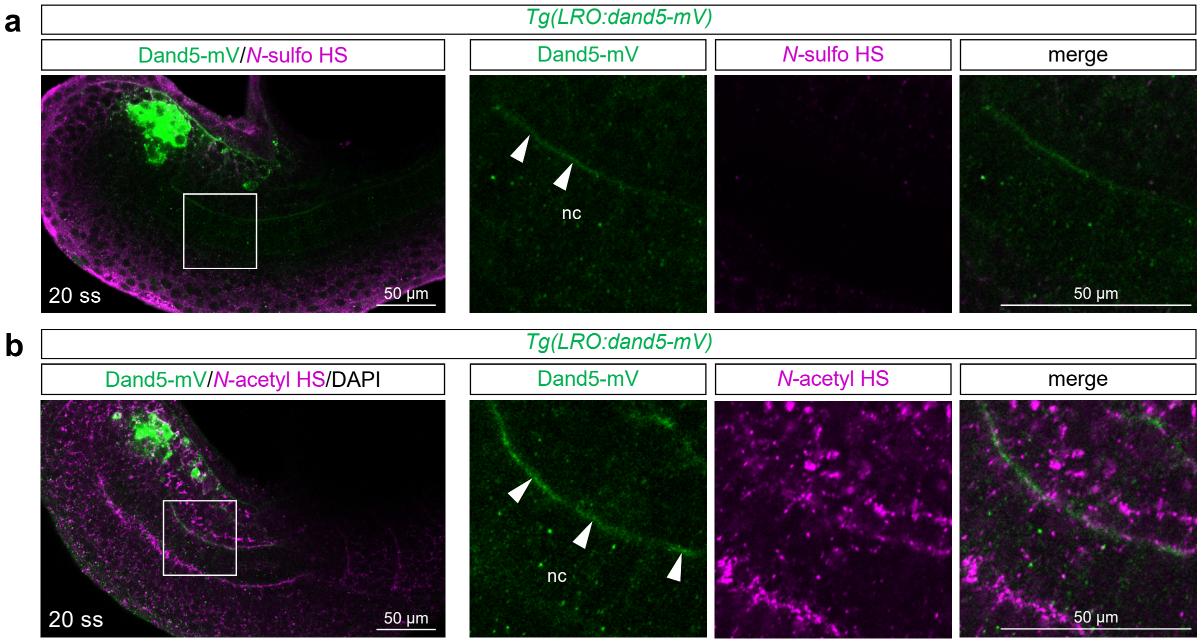

### Supplementary Figure 7

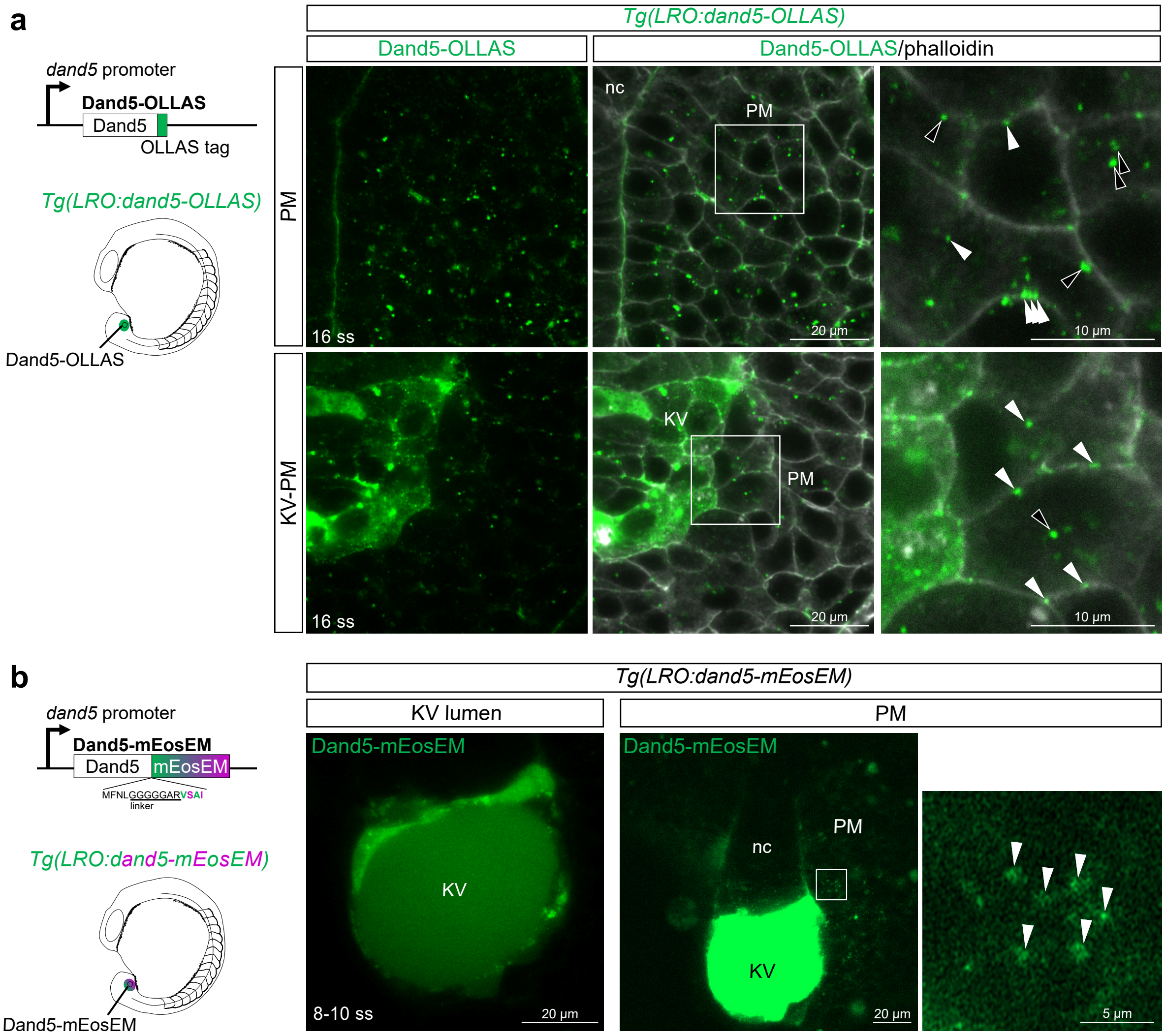

### Supplementary Figure 8

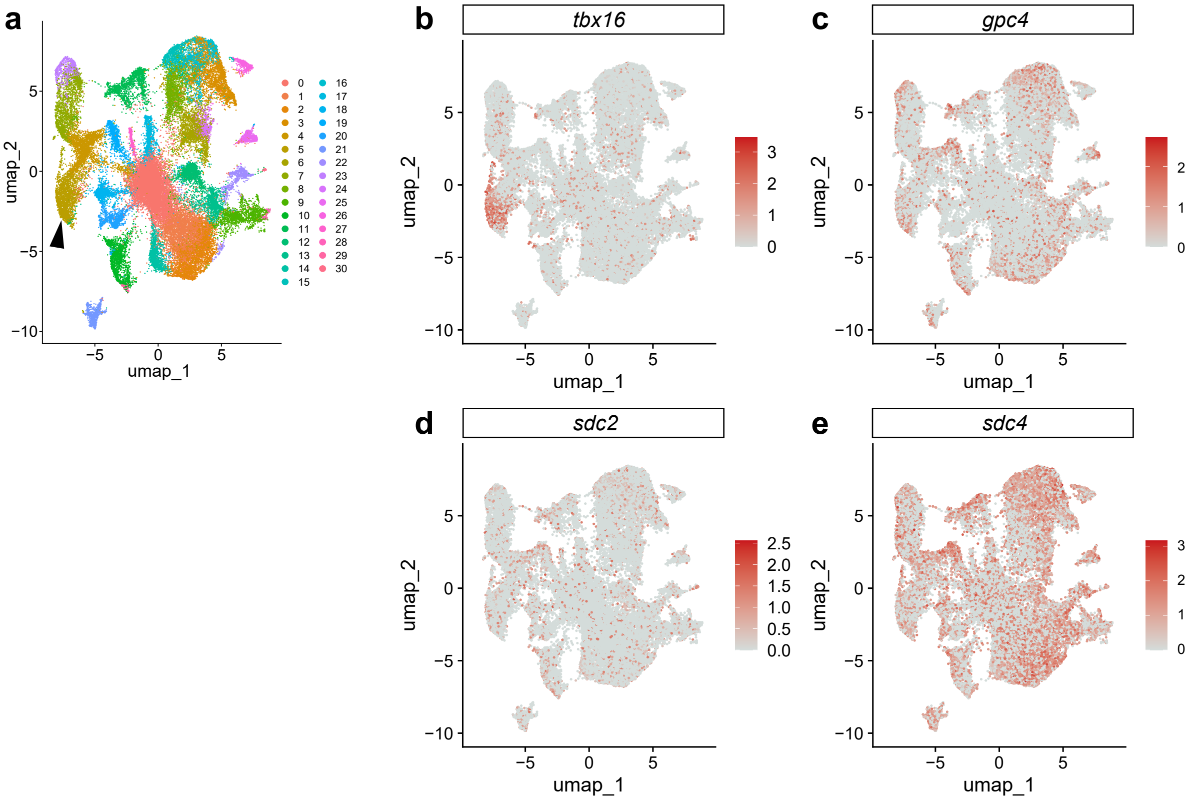

### Supplementary Figure 9

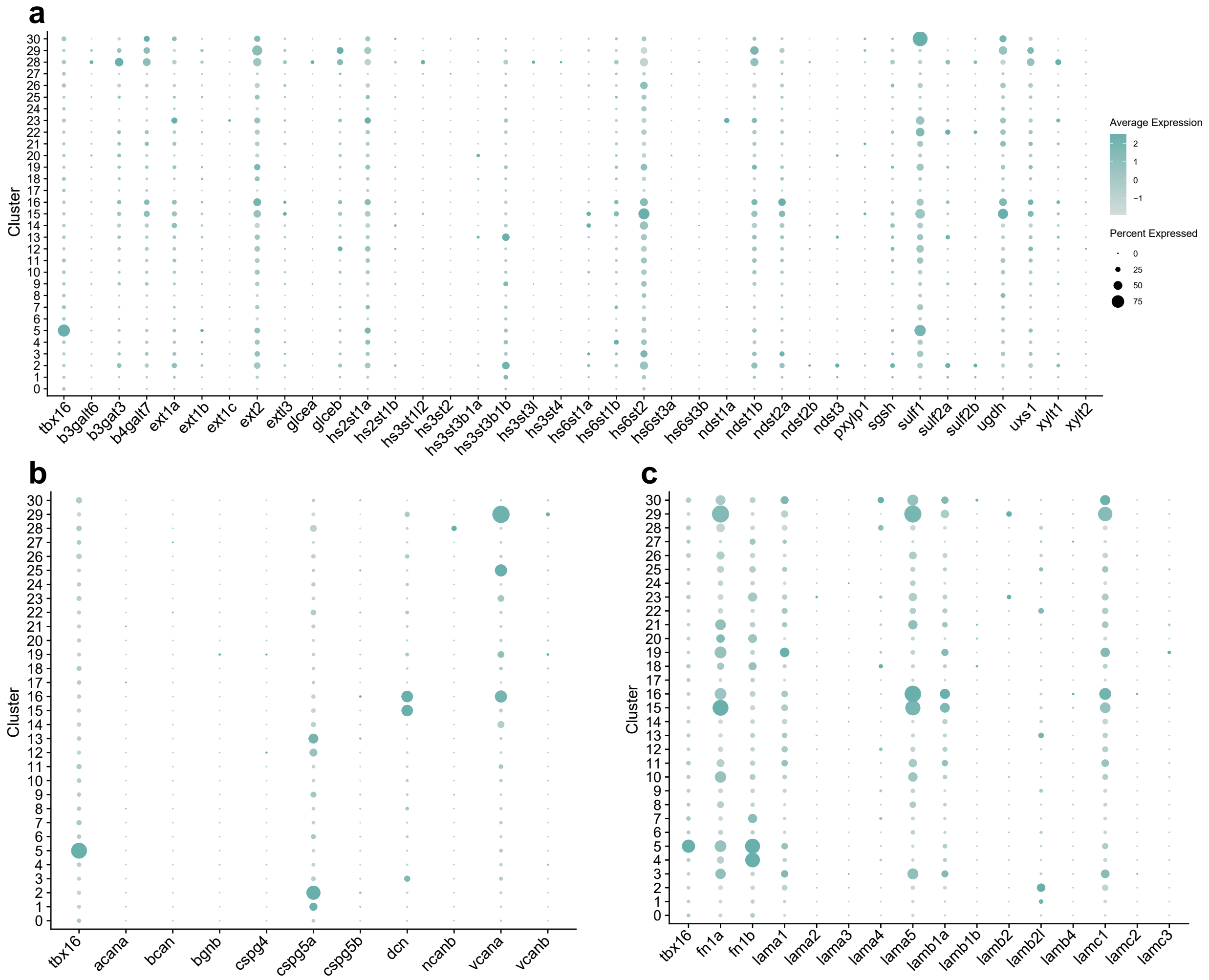

### Supplementary Figure 10

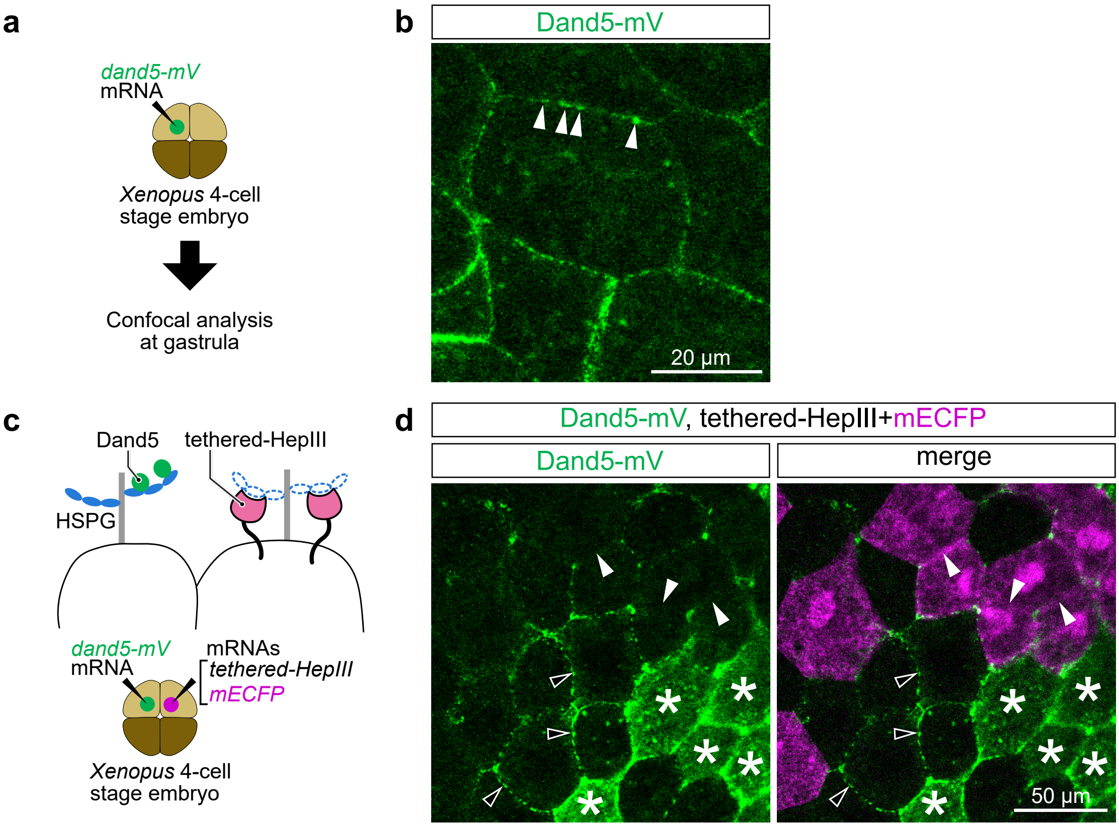

### Supplementary Figure 11

**a**

>Dand5
MTFQVGFFVLLSVTTIGAFPRNAFQREFHRHVAKDFESSGNGPDEPVRGSVRIVKLNPHFLRR
AAVSHVPFRNSPSRGAFPAFLALGRPGPAILTHSKPAPQVSSSADRRKQGLEMWKKVVHKSER
KKEAVALRINPKDMNKQSCAAVPFTQRITEEGCETVTVHNNLCYGQCSSMFVPSSGGSHGQQK
AQCMRCGPSRARSVLLHLRRGSEVRERRVLIVEECKCETSSEEAKVQNTDMFNL*

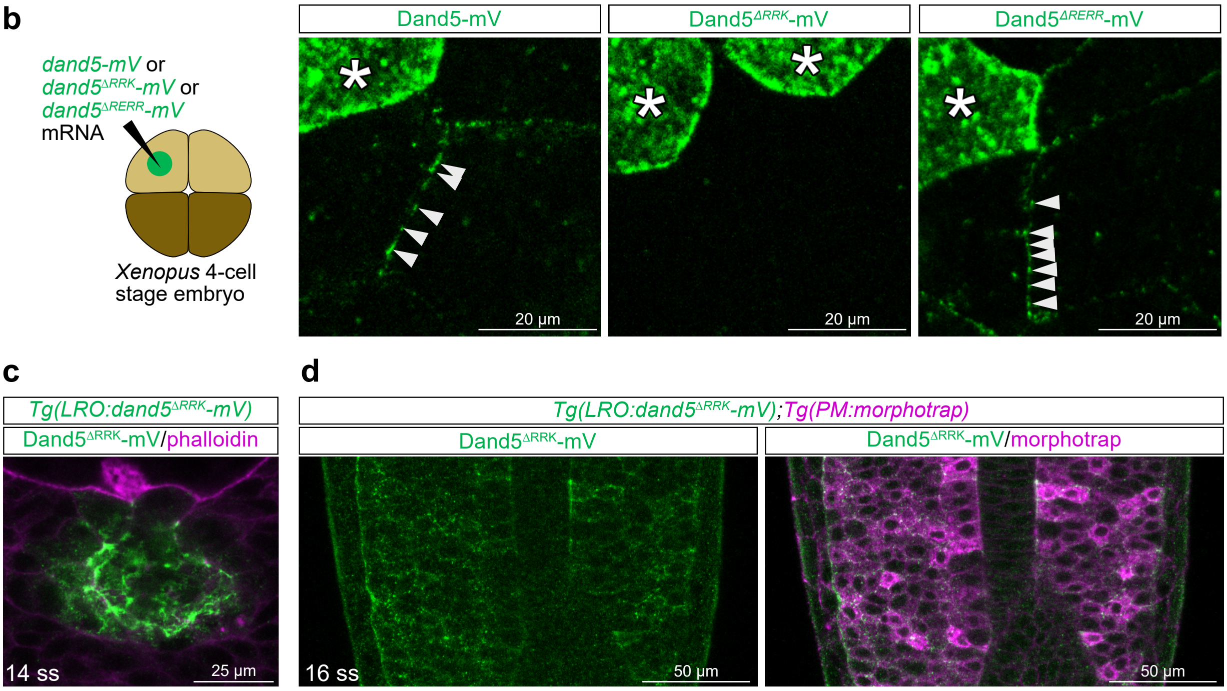

### Supplementary Figure 12

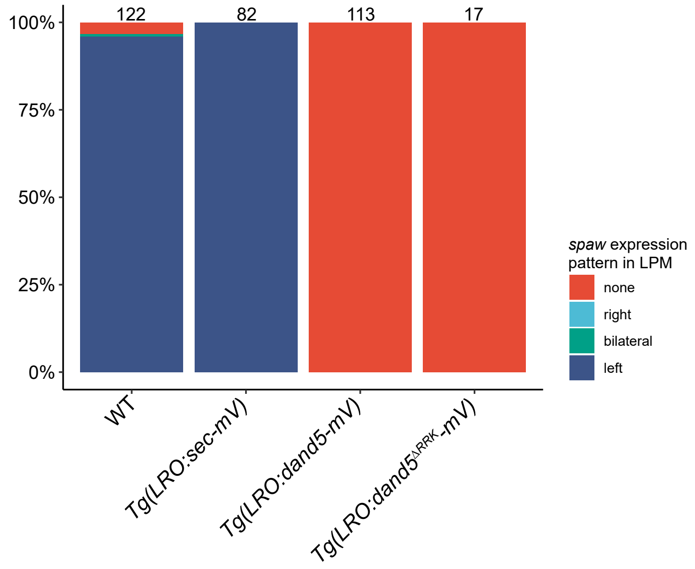

### Supplementary Figure 13

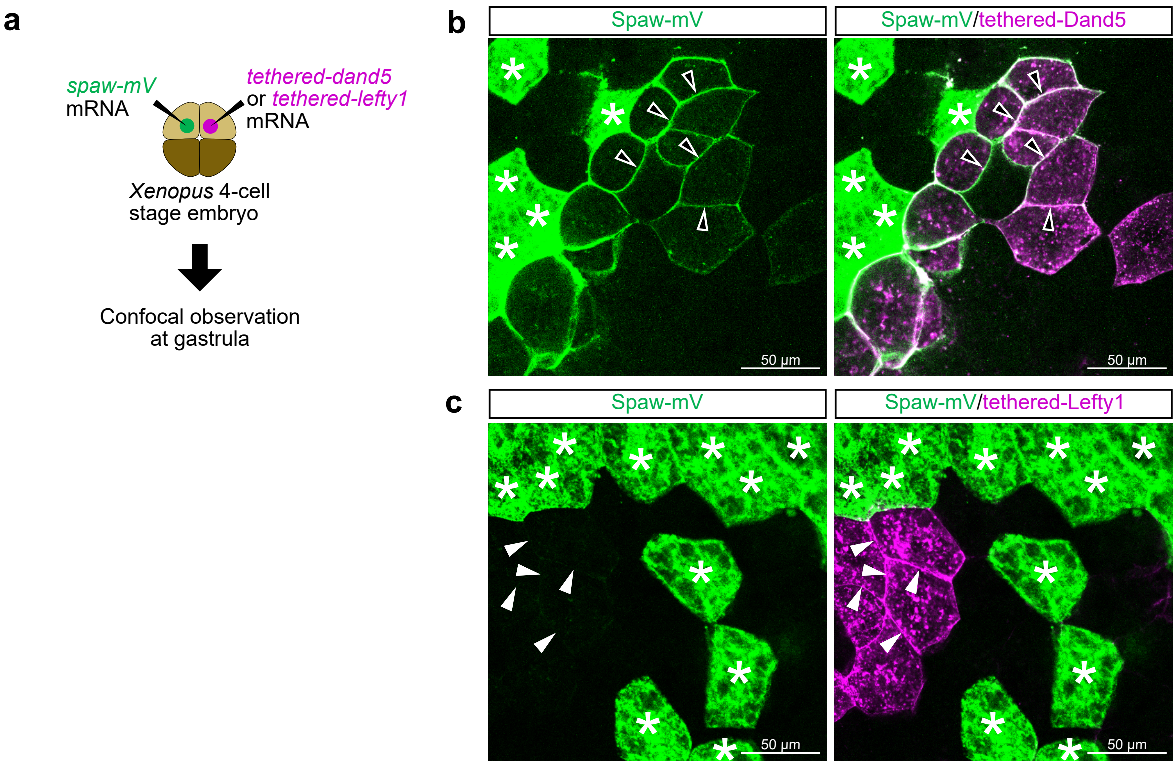

### Supplementary Figure 14

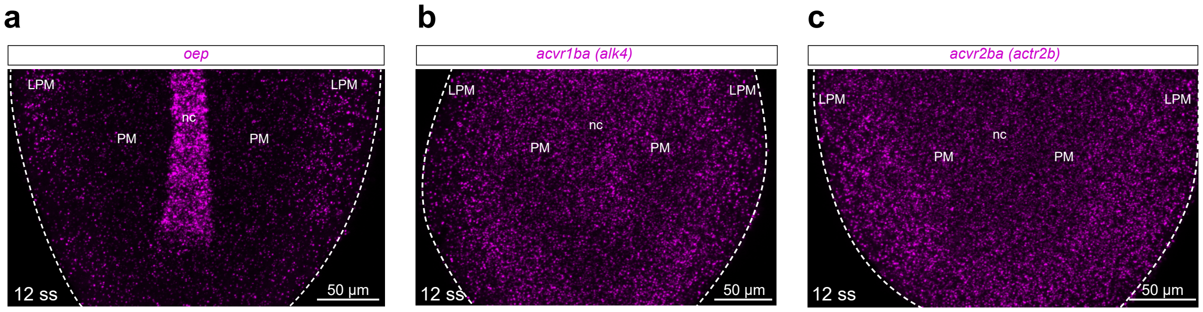
